# Network-aware self-supervised learning enables high-content phenotypic screening for genetic modifiers of neuronal activity dynamics

**DOI:** 10.1101/2025.02.04.636489

**Authors:** Parker Grosjean, Kaivalya Shevade, Cuong Nguyen, Sarah Ancheta, Karl Mader, Ivan Franco, Seok-Jin Heo, Greyson Lewis, Dehua Zhao, Bhairavi Tolani, Steven Boggess, Angelique Di Domenico, Erik Ullian, Shawn Shafer, Adam Litterman, Laralynne Przybyla, Michael J. Keiser, Jamie Ifkovits, Adam Yala, Martin Kampmann

## Abstract

High-throughput phenotypic screening has historically relied on manually selected features, limiting our ability to capture complex cellular processes, particularly neuronal activity dynamics. While recent advances in self-supervised learning have revolutionized the ability to study cellular morphology and transcriptomics, dynamic cellular processes have remained challenging to phenotypically profile. To address this limitation, we developed Plexus, a self-supervised model specifically designed to capture and quantify network-level neuronal activity dynamics. Unlike existing phenotyping tools that focus on static readouts, Plexus leverages a network-level cell encoding method, which enables it to efficiently encode dynamic neuronal activity data into rich representational embeddings. In turn, Plexus achieves state of the art performance in detecting phenotypic changes in neuronal activity. We validated Plexus using a comprehensive GCaMP6m simulation framework and demonstrated its enhanced ability to classify distinct neuronal activity phenotypes compared to traditional signal-processing approaches. To enable practical application, we integrated Plexus with a scalable experimental system utilizing human iPSC-derived neurons equipped with the GCaMP6m calcium indicator and CRISPR interference machinery. This integrated platform successfully identified nearly seventeen times as many distinct phenotypic changes in response to genetic perturbations compared to conventional signal processing methods, as demonstrated in a 52-gene CRISPRi screen across multiple iPSC lines. Using this framework, we identified potential genetic modifiers of aberrant neuronal activity in frontotemporal dementia, illustrating its utility for understanding complex neurological disorders.

## Introduction

Advances in high-content microscopy have enabled a new era of cellular phenotyping. Combined with genetic and small molecule perturbations, high-throughput phenotyping promises to provide insight into how genes and chemical matter drive or modify cellular processes^1–8^. However, profiling dynamic cellular processes, such as neuronal activity, remains a grand challenge; as a result, it remains difficult to identify the genes underpinning complex neuronal diseases, such as epilepsy and neurodegeneration. Our study aims to enable phenotypic screening in networks of neurons, opening doors for biological discovery. Our contributions to address this end are threefold – we (1) engineered an experimental cellular system to study neuronal activity dynamics *in vitro*, (2) developed a self-supervised model, Plexus, to identify biologically distinct phenotypes, and (3) we established a CRISPR interference (CRISPRi) screening platform to uncover genes that regulate neuronal activity. Building on these foundations, we were able to identify genes that may be involved in aberrant dysregulated neuronal activity in a neurodegenerative disease, frontotemporal dementia.

Current experimental approaches to quantifying neuronal activity, such as patch-clamp electrophysiology^9^ or multi-electrode arrays^8^, while powerful for detailed characterization, are limited in throughput and resolution, respectively. More recent approaches leveraging genetically encoded calcium^10,11^ and voltage sensors^12^ have shown promise in increasing screening throughput for small molecule drug development. However, these approaches have solely focused on intrinsic membrane excitability. Symptoms of neurological diseases often manifest as dysregulated network-level (synaptic) activity, so experimentally modeling and computationally quantifying both homeostatic and aberrant network-level phenotypes *in vitro* is crucial when screening for protein targets with disease-modifying effects. To address this, we established a human induced pluripotent stem cell (hiPSC)-derived neuron and astrocyte co-culture model equipped with the GCaMP6m calcium indicator^13^, enabling scalable measurements via microscopy of both single-cell and network activity dynamics. We also engineered our system to incorporate CRISPR interference (CRISPRi) machinery, enabling the targeted knockdown of gene expression. This provides a tractable system for studying neuronal activity and the genes involved in its regulation.

Phenotypic profiling (quantitatively measuring biological features in high throughput) has a rich history, with early examples relying on hand-selecting a single feature believed to be useful in a cellular process of interest^14^. However, many critical cellular states and processes are inherently high-dimensional and are not amenable to manual feature engineering. This has led to the rapid adoption of self-supervised learning (SSL)-based profiling methods, enabling biologists to learn features directly from the underlying biological data^15–18^. However, these tools have focused primarily on morphometric and transcriptomic readouts, and they are not designed to provide insight into dynamic cellular processes, such as neuronal activity. To model neuronal activity dynamics and uncover the genes that regulate them, we designed an SSL-based model, Plexus. Through both modeling and pre-training innovations, Plexus is tailored to capture network activity dynamics. Across a wide range of experiments, we demonstrate that Plexus outperforms both single-cell level SSL and state-of-the-art signal processing-based approaches^10^.

We developed a GCaMP6m simulation framework to benchmark Plexus’ performance against previously existing phenotyping methods. By parametrically modeling GCaMP6m signals with different levels of synaptic strength, negative regulation, and intrinsic excitability, we demonstrate via linear probing (training and evaluating a linear model on the embeddings generated from the frozen model) that Plexus provides state of the art classification accuracy for simulated neuronal activity phenotypes generated with distinct parameters. Next, to validate our computational and experimental platform together, we tested the ability of Plexus to identify activity phenotypes driven by chemical perturbations in our experimental system. Using linear probing we demonstrate Plexus’ superior performance in capturing neuronal activity phenotypes in experimental data compared to prior methods.

Building on Plexus and our experimental system, we developed a platform for running arrayed CRISPRi-based genetic perturbation screens. To validate our platform, we performed a CRISPRi screen targeting 52 genes. We performed this screen in two hiPSC lines with different genetic backgrounds, demonstrating that the platform is robust across distinct iPSC cell lines. Plexus detected on average 22.5 times more phenotypic changes in response to gene perturbation compared to traditional signal processing feature-based approaches, when using a mean two-fold AUROC of 0.8 as a threshold for a distinct phenotype. To validate that the gene perturbation-driven phenotypes we uncover are biologically meaningful, we focused on the phenotype driven by the knockdown of the gene *KCNQ2*. Using an interpretability analysis, we dissected the learned embeddings through the lens of manually defined features to show a phenotype consistent with known physiology in both iPSC-derived cell lines.

Building on these foundations, we shifted focus to screening for genes that may play a role in frontotemporal dementia, a neurodegenerative disease. We extended our CRISPRi screening approach to a model of aberrant neuronal activity associated with frontotemporal dementia and used Plexus to identify genes that drive phenotypic changes toward healthy network states. Altogether, this study introduces an experimental and computational framework to facilitate the discovery of disease-specific modifiers of neuronal activity.

## Results

### Enabling high-content phenotyping of neuronal dynamics *in vitro* using a human iPSC-derived neuron and astrocyte co-culture system

To establish a readout of neuronal activity dynamics, we sought to generate a system that recapitulates key components of neuronal activity – excitatory synaptic activity and spontaneous depolarization events. To accomplish this, we focused on developing a fully human, induced pluripotent stem cell (hiPSC)-derived co-culture system of astrocytes and excitatory neurons. To rapidly generate homogenous excitatory cortical-like neurons, we overexpress the transcription factor Neurogenin 2 (NGN2) from a safe-harbor locus via induction with doxycycline^19^. We will hereafter refer to these cells as iNeurons. We engineered these iNeurons to express the CRISPR interference (CRISPRi) machinery^20^ and constitutively active GCaMP6m^13^, enabling the targeted knockdown of genes and measurement of calcium as a secondary readout of neuronal activity, respectively (**Figure 1A**). To ensure biologically relevant neuronal activity phenotypes, we adapted a protocol^21^ to generate mature CD49f+ astrocytes from hiPSCs using a line lacking the aforementioned CRISPRi machinery or GCaMP6m reporter system (**Figure 1B**). We incorporated astrocytes into our model system, as they are essential to maintaining physiologically relevant levels of neuronal activity, which we confirmed using a multi-electrode array (**Extended Data Figure 1**). To showcase the efficacy of the *in vitro* model system, we cultured our iNeurons and astrocytes and imaged their protein expression and calcium fluctuations at days 14 and 21 (**Figure 1C**). The neuronal activity was measured on day 14 because this was found to be the earliest time point where all assayed neuronal cultures were active (**Extended Data Figure 1**). We additionally imaged at day 21 for longitudinal monitoring of activity during differentiation. Imaging the astrocytes, we demonstrate that they are positive for the canonical astrocyte markers S100B and GFAP in co-culture with the GCaMP6m iNeurons (**Figure 1D**).

**Figure 1:**
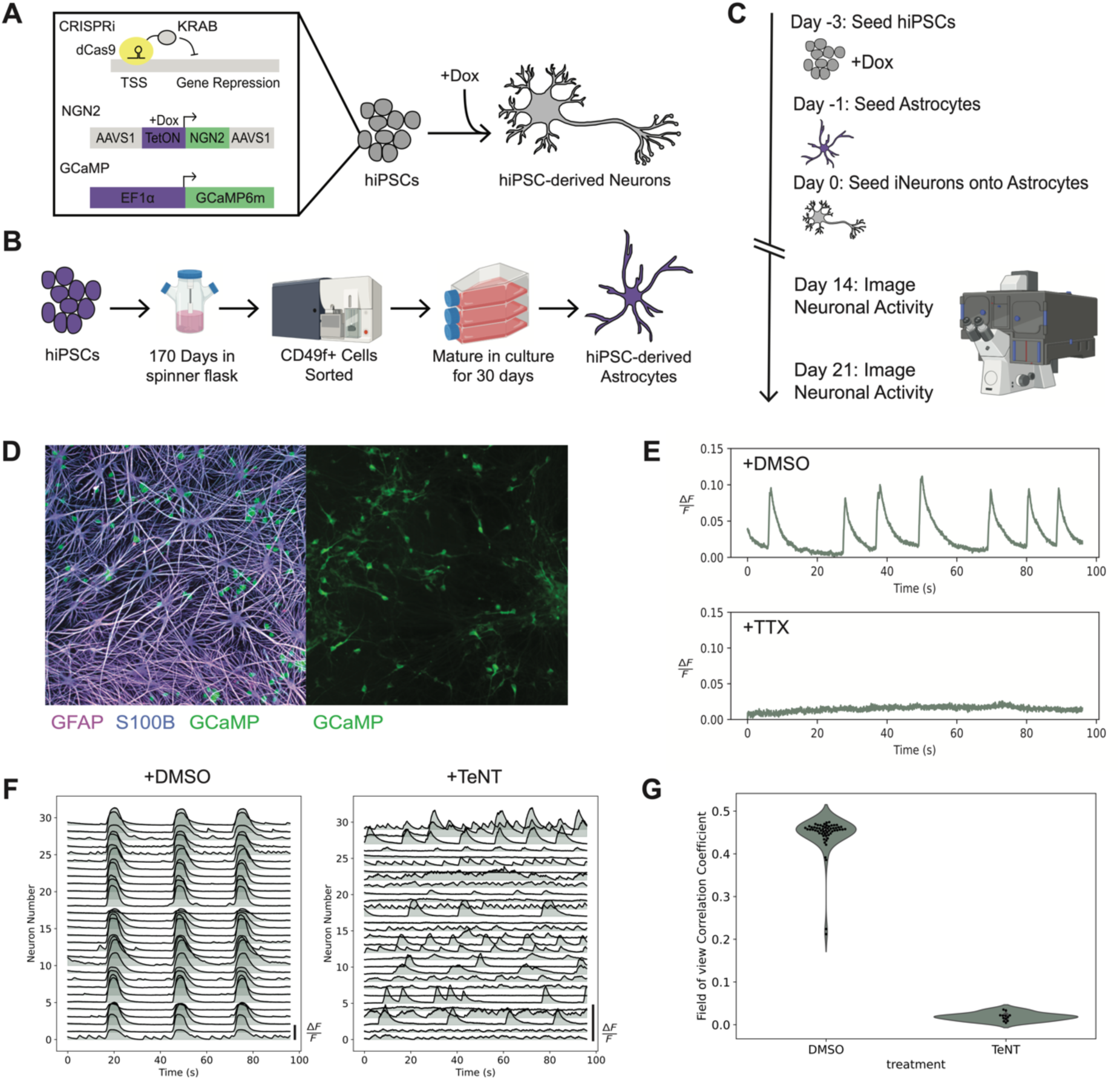
Neuronal activity assay overview. (A) Schematic of the inducible neuron system containing CRISPRi machinery, GCaMP6m, and a doxycycline-inducible NGN2 expression cassette. (B) Overview of organoid-derived astrocyte generation. (C) Schematic of the neuron-astrocyte co-culture model’s plating, differentiation, and imaging. (D) Immunostaining of canonical astrocyte markers (S100B, GFAP) and an example image of the genetically encoded calcium indicator GCaMP6m. (E) Spontaneous neuronal activity is silenced upon treatment with tetrodotoxin (TTX). (F) Representative calcium fluctuations in DIV 21 neurons exhibit basal synaptic activity and abrogation upon treatment with tetanus toxin (TeNT). (G) The field of view correlation coefficient in TeNT-treated (n=6) neuronal cultures is decreased compared to DMSO-treated (n=29) cultures.

To confirm that the calcium fluctuations observed via GCaMP6m were due to full membrane depolarization events, the neuronal cultures were treated with tetrodotoxin (TTX). This potent sodium channel blocker silenced the calcium oscillations in the culture, showing that the iNeurons are spontaneously active in our cultures (**Figure 1E**). In addition to spontaneous depolarization events, network-level phenotypes are driven by synaptic transmission. To ensure that the cultures were synaptically active, we imaged the co-cultures and observed spontaneous synchronous synaptic activity via GCaMP6m, which was ablated upon treatment with tetanus toxin (TeNT), a neurotoxin that inhibits the release of presynaptic vesicles by disrupting the SNARE complex ^23^ (**Figure 1F**). The abrogation of synaptic activity was quantified by measuring the drop in the correlation coefficient of neurons firing synchronously in the same field of view (**Figure 1G**). The *in vitro* astrocyte and neuron co-culture model demonstrates spontaneous depolarization events and synaptic transmission, providing a tractable system for studying the effect of gene expression and neuroactive compound perturbations.

### Plexus: Network-aware self-supervised learning for neuronal activity phenotypic profiling

Homeostatic neuronal activity is a function of intrinsic excitability, synaptic excitability, and network-level properties. Dysregulated neuronal activity can arise from changes to any of these properties. The goal of phenotypic profiling is to capture all aspects of neuronal activity within concise computational representations, termed latent vectors, such that the vectors arising from distinct neuronal phenotypes are linearly separable. Distilling neuronal activity into a well-organized latent space would empower researchers to reason over complex biological perturbations using simple vector arithmetic. To this end, we introduce Plexus, our computational framework for neuronal activity phenotypic profiling. Plexus introduces key innovations in neural network architectures and self-supervised learning to model networks of neurons efficiently.

To capture arbitrary dependencies between the spike trains of neurons within a network, we propose to adapt Transformers^22^. We consider each neuron’s spike train as a sequence of small event windows (i.e., tokens); we can then model network activity as an extended sequence of spike trains, where each token derives from a particular cell and activity time point. We can provide the Transformer with knowledge of each token’s source cell and time point using positional embeddings. While this simple framework allows us to capture arbitrary dependencies between input tokens, efficiently modeling network activity requires accounting for the inherent symmetries within our modality.

Neurons in networks have no canonical ordering or position; moreover, randomly assigning neurons positional embeddings can complicate learning dynamics, as each permutation of neurons would result in a distinct embedding. To overcome this challenge, we introduce a cell identification embedding layer. We propose to induce a global ordering of cells within each subnetwork using the average L2 norm within each cell’s input signal; this subnetwork-specific ranking is then used to index into learned positional embeddings to identify each cell (**Figure 2A**). Thus, for each subnetwork, every cell, regardless of permutation (a cell could be the first n tokens or the last n tokens in the input sequence), will always receive the same learnable cell identification embedding. We chose this simple identification scheme (indexing into discrete learnable embeddings) over other permutation-invariant approaches such as using a single cell-specific feature, as it allows us to generate a canonical representation for each subnetwork of cells, enabling efficient learning.

**Figure 2:**
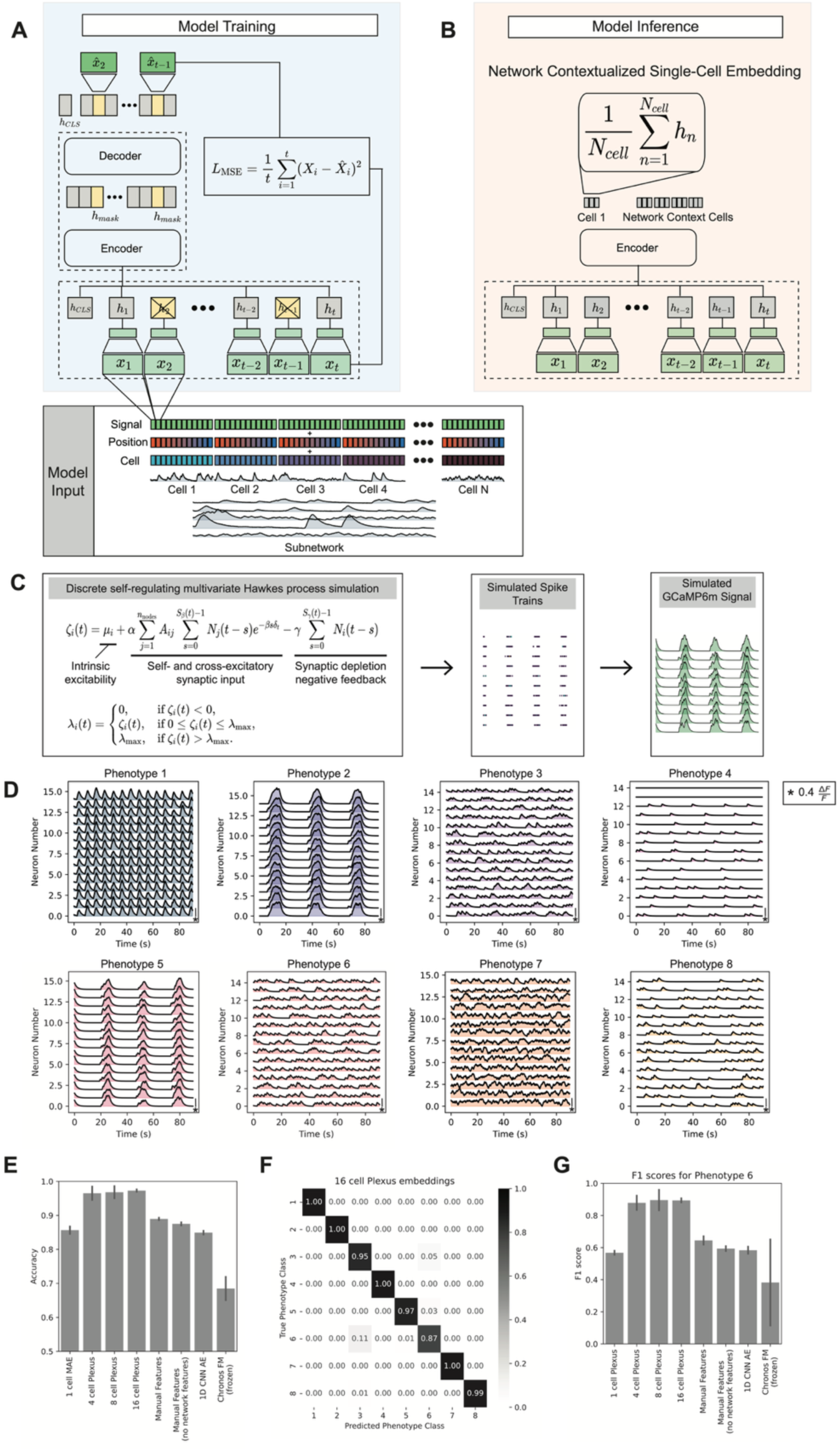
Network-aware self-supervised masked autoencoder and validation on simulated data. (A) The self-supervised model is trained on sampled subnetworks of activity traces chunked into time windows for input into the model. Each time window is then linearly projected to tokens. Then, a portion of all the tokens is randomly masked, and only the unmasked tokens are fed into the encoder of the Plexus to generate the latent embeddings. The masked tokens are added back with the latent embeddings and passed to the decoder to use the latent information in the unmasked tokens to reconstruct the masked time windows using a mean-squared error loss term. (B) To generate network contextualized single-cell embeddings, all tokens are left unmasked and passed through the Plexus encoder with the model weights frozen. Single-cell embeddings are generated by mean pooling over the latent embeddings of all tokens belonging to each cell in the subnetwork. (C) The simulation framework for GCaMP data uses a self-regulating multi-variate Hawkes process to generate neuronal spiking events, which were converted to estimated calcium concentration fluorescence signal using the autoregressive model with 2 degrees of freedom developed by Song et al.^27^. (D) Representative calcium traces from the eight distinct neuronal activity phenotypes were generated using the simulation framework with varied parameters. (E) Linear probing accuracies for the eight-class classification task using a three-fold cross-validation split (n=14440 for each class; Error bars are standard deviation) demonstrate Plexus embeddings outperform manual features and other ML-derived baselines in phenotypic classification. (F) Confusion matrix for the 16-cell Plexus embedding linear probing classification task. (G) The F1 scores for the linear probing classification of phenotype 6 demonstrate increased precision and recall of the Plexus model for difficult phenotype classification tasks compared to other manual feature and ML-based baselines.

To obtain linearly separable representations using Plexus, we propose a self-supervised learning algorithm. We randomly sample subnetworks of neurons and then randomly mask a fraction of the resulting tokens. In doing so, we randomly drop temporal chunks of activity across cells within the subnetwork. We then train Plexus to recover the masked tokens across the subnetwork. Specifically, we input unmasked tokens into our Transformer encoder to generate network-contextualized representations; we then pass these contextualized tokens and the masked tokens into a transformer decoder to predict the masked activity **(Extended Data Figure 2)**. This self-supervised method is an extension of Masked Auto Encoders^23^ (MAE) that explicitly forces the underlying model to incorporate network information. After self-supervised training, we can use Plexus to generate network-aware single-cell representations (**Figure 2B**). We pass the entire neuronal network of activity traces into our Transformer encoder and then leverage mean pooling across the tokens of each cell to obtain single-cell summary embeddings. This approach allows us to quantify cellular heterogeneity while capturing synaptic activity from the network.

### Validating Plexus: Simulation-based experiments demonstrate the identification of distinct neuronal activity phenotypes

To characterize how well profiling approaches capture distinct and diverse neural activity phenotypes, we developed a simulation-based benchmark. By controlling discrete aspects of neuronal activity in a multivariate Hawkes process, we can generate distinct phenotypes with varying difficulty levels; this explicit control offers a rich resource to understand sources of model failure, which is difficult to obtain with in-vitro experimental data where no “ground truth” is available. In this framework, we compared Plexus to a single-cell MAE pretrained Transformer, and traditional manual features.

Our simulations leverage mathematical point processes to model neuronal depolarization events or spikes. Instead of leveraging a Poisson process which assumes that events occur independently and uniformly in time, we leverage a self-regulating multi-variate Hawkes process to account for the synaptic feedback and self-excitatory processes observed in neuronal activity. Instead of having a univariate scalar intensity function like a Poisson process, we leverage a multivariate time-dependent intensity function, building on the model presented in Goncalves et al.^24^ The intensity function includes terms to capture: (1) the intrinsic excitability of a neuron, (2) the cross- and self-excitation (synaptic neurotransmitter reuptake rate, short-term potentiation, number of synapses, and synaptic release probability), and (3) negative self-regulation of firing (depolarization blocks and short-term depression). To model distinct network spiking behavior, we generated Watts-Strogatz networks^25^ with a fixed number of nodes (neurons). We then modeled each neuron within the network by discretely sampling from the Poisson distribution with the intensity function evaluated at each timestep (**Methods**). We generated eight distinct neuronal activity spike event phenotypes (**Extended Data Table 1**); we converted each spiking phenotype into simulated GCaMP6m signals following the method developed by Song et al.^26^ (**Figure 2C**) which resulted in calcium activity (**Figure 2D**) with different synaptic strengths, temporal feedback windows, and basal excitatory activity. We chose these eight phenotypes, as we found they were sufficient to reflect the diversity of phenotypes observed in our *in vitro* co-culture model (**Extended Data Figure 3**).

Given these simulated data, we trained a single-cell MAE pretrained Transformer (1-cell) and our Plexus model with varying subnetwork sizes (4-cell, 8-cell, and 16-cell) with a 50% masking ratio for 1000 epochs and extracted embeddings (**Methods**). We also calculated manual features based on signal processing techniques as a baseline phenotyping method (**Methods**). To determine model performance, we performed linear probing for the 8-class phenotype classification task with a 3-fold well-aware cross-validation. Well-aware splits are used to ensure that well-specific synchronicity patterns are not used as a shortcut for classification. For this linear probing task, higher classification accuracy denotes better performance. We found that a single-cell MAE performs worse (0.8568 ± 0.0093 accuracy), a 1D CNN autoencoder, and a pre-trained time-series foundation model^27^ all performed worse than manually engineered features (0.8901 ± 0.0025). However, upon the addition of any contextualizing cells from the subnetwork, the linear probing accuracy increases (4-cell: 0.9641 ± 0.0164, 8-cell: 0.9691 ± 0.0155, 16-cell: 0.9729 ± 0.0027) (**Figure 2E**). The 4-cell, 8-cell, and 16-cell Plexus embeddings had comparable linear probing accuracies, suggesting that providing any number of contextualizing cells boosts the ability to discern the simulated phenotypes from one another.

The confusion matrix of the 16-cell Plexus demonstrates that while most class accuracies were close to the overall performance of the model, one phenotype – phenotype 6 – had a lower accuracy (**Figure 2F**). This lower accuracy is driven primarily by confusion with phenotype 3, which is consistent with the only difference in the simulation parameters being either no synaptic input in the case of phenotype 3 or weak synaptic input in phenotype 6 (**Extended Data Table 1**). To investigate this further, we calculated the macro F1 score for phenotype 6 for every set of embeddings and manual features (**Figure 2G**). While the linear probing accuracy on the 16-cell Plexus embeddings for this phenotype is the lowest, this demonstrates the higher sensitivity of the Plexus embeddings to changes in the neuronal activity driven by synaptic input compared to manual features both with and without features that measure correlation with other cells in the network.

### Validating Plexus and Co-Culture System: Neuroactive stimulation experiments

After demonstrating the ability of the Plexus model embeddings to capture the differences in simulated neuronal activity phenotypes, we investigated its performance in capturing experimentally measured neuronal activity dynamics driven by treatment with known activity modulators (**Figure 3A**). These experiments validate both the experimental platform and profiling algorithm. We used tools that are known to decrease and ablate synaptic activity (2 mM Mg^2+^, TeNT) as well as increase intrinsic excitability and synaptic activity (2 mM Ca^2+^) to investigate the model performance. In addition, we used TTX to ablate intrinsic excitability via its potent ability to block voltage-gated sodium channels. Each treatment was added to a set of wells in a 384-well plate, such that there were multiple biological replicates for every treatment. This enabled us to quantify the generalizability of phenotypes across wells.

**Figure 3:**
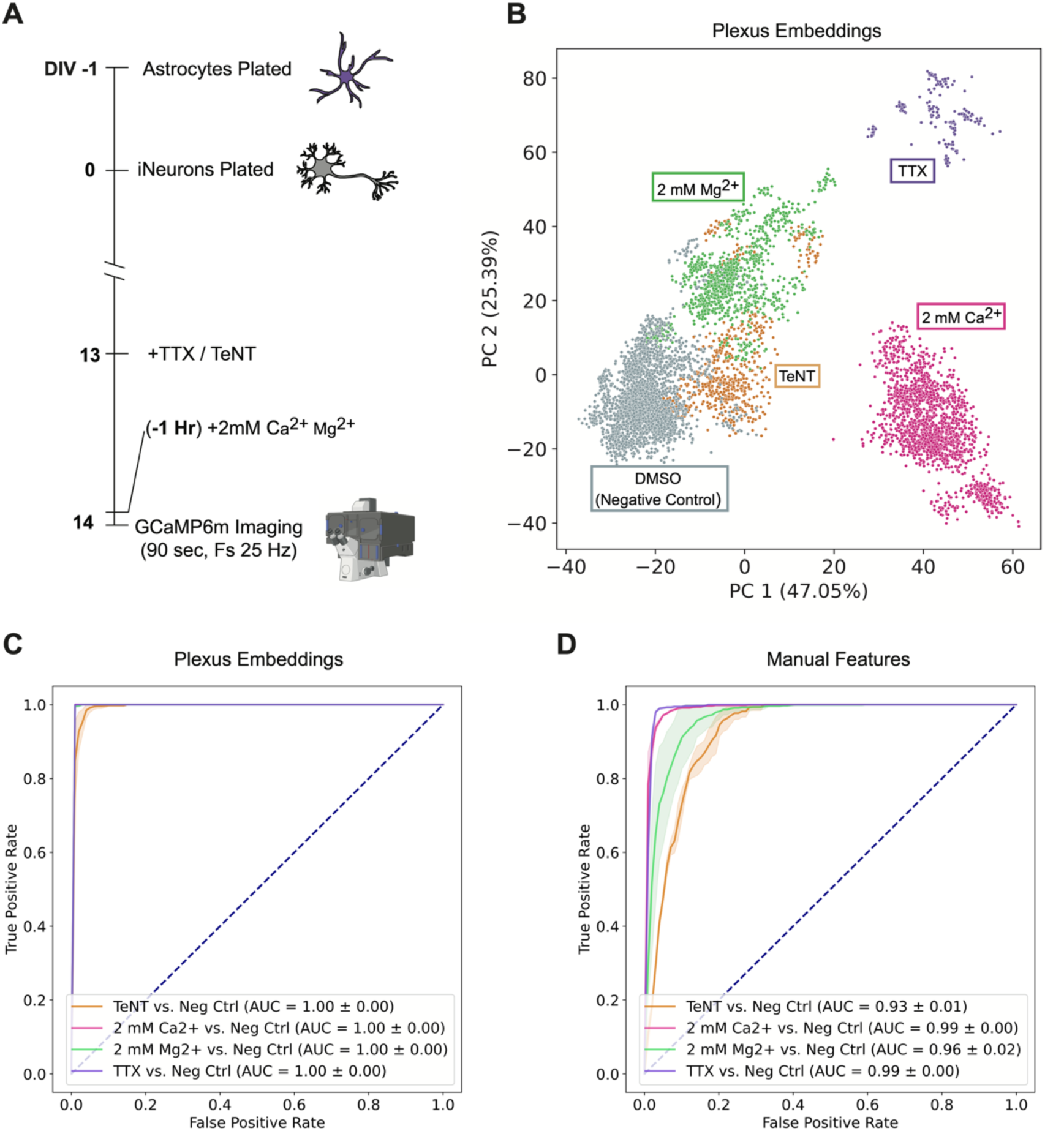
Plexus captures separable phenotypic profiles of neuroactive treatments *in vitro* (A) We performed a neuroactive treatment assay with GCaMP6m iNeurons, with TTX and TeNT treatment 24 hours before imaging and 2 mM Mg^2+^ and 2 mM Ca^2+^ added 1 hour before imaging. (B) Projection of the single-cell network contextualized embeddings onto the first two principal components shows the separation between distinct neuroactive treatment phenotypes. (C) Receiver operator curves for elastic-net binary classifiers trained between negative control and neuroactive treatments (TeNT n= 688, TTX n=1773, Negative Control n=3723, 2 mM Mg2+ n=1741, 2 mM Ca2+ n=1942). (D) Receiver operator curves for elastic-net binary classifiers trained on engineered features for negative control treatment versus neuroactive treatments; same sample sizes as (C).

We trained Plexus for 1000 epochs on the activity traces (**Methods**). Plexus embeddings capture differences in the activity phenotypes, as shown by the low-dimensional projection of the embeddings onto the first two principal components (**Figure 3B**).

We assessed the ability to distinguish between activity phenotypes based on the Plexus embeddings by training elastic-net binary classifier models to separate the negative control cells from those treated with neuroactive agents. We used a three-fold cross-validation split with well-wise stratification, so that cells from the same well were always in the same fold, to verify that model performance is not based on well-specific information. The area under the receiver operating curve (AUROC) for the elastic-net binary classifier models demonstrated the linear separability of the negative control and treated embeddings (2 mM Mg^2+^ = 1.00, TeNT = 0.98, 2 mM Ca^2+^ = 1.00, TTX = 1.00; **Figure 3C**). To ensure that spurious correlations and shortcut learning were not driving the cross-split performance of the models, we trained an elastic-net binary classifier to distinguish between random well-splits of the negative control and observed roughly random classification performance (AUROC 0.52). We also compared the separability of the activity phenotypes using manual features similar to those used by Boivin et al.^10^(**Methods**). Linear probing of the manual feature vectors derived from signal processing reduced average AUROCs on the treatment separation task (2 mM Mg^2+^ = 0.96, TeNT = 0.93, 2 mM Ca^2+^ = 0.99, TTX = 0.99; **Figure 3D**).

### Enabling CRISPR interference screening for uncovering genetic modifiers of neuronal activity dynamics

With the ability to distinguish between distinct *chemical-matter induced* neuronal activity phenotypes, we focused on leveraging this approach to understand genetic etiologies of altered neuronal activity.

To build our CRISPRi screening platform, we developed a library of dual-guide RNA constructs targeting 52 genes with putative roles in neuronal activity or related neurological conditions, in addition to non-targeting negative controls that do not target genomic sequence, controlling for the effect of lentiviral transduction (**Supplementary Table 2**). The library consisted of one dual guide construct targeting each gene. Dual-guide vectors were chosen as they have been shown to increase the on-target knockdown efficiency and the power of CRISPRi screening^28^. To this end, we generated an arrayed plasmid library of the dual-guide vectors and subsequently generated an arrayed lentiviral library containing the dual-guide constructs (**Methods**). To ensure homogenous iNeuron populations, pre-differentiation of the inducible iNeurons with doxycycline was performed in large single batches and frozen in aliquots for screening. The mature iPSC-derived astrocytes were plated at D-1 of the screen into 384-well plates, followed by the robotic addition of the arrayed lentivirus on D0 (**Figure 4A**). The aliquots of homogenous pre-differentiated iNeurons were added to the 384-well plates for reverse transfection, such that each well had exactly one dual-guide construct delivered. These cultures were then differentiated for 14 days and imaged at 25 Hz for 90 seconds per field of view, with two fields of view per well. The dual-guide constructs contain a nuclear-localized blue fluorescent protein (BFP) element, enabling us to determine which iNeurons had been successfully transduced (**Figure 4B**). We determined that the robotic reverse transduction of the iNeurons enabled a consistent transduction efficiency of ∼60% across all wells (**Extended Data Figure 4)**. In total, each dual-guide lentiviral vector was delivered to 8 wells across two different genetic backgrounds (cell line 1: WTC11^29^, cell line 2: Patient^30^), spanning eight total 384-well plates. This resulted in 16 field-of-view videos per dual-guide vector or gene knockdown.

**Figure 4:**
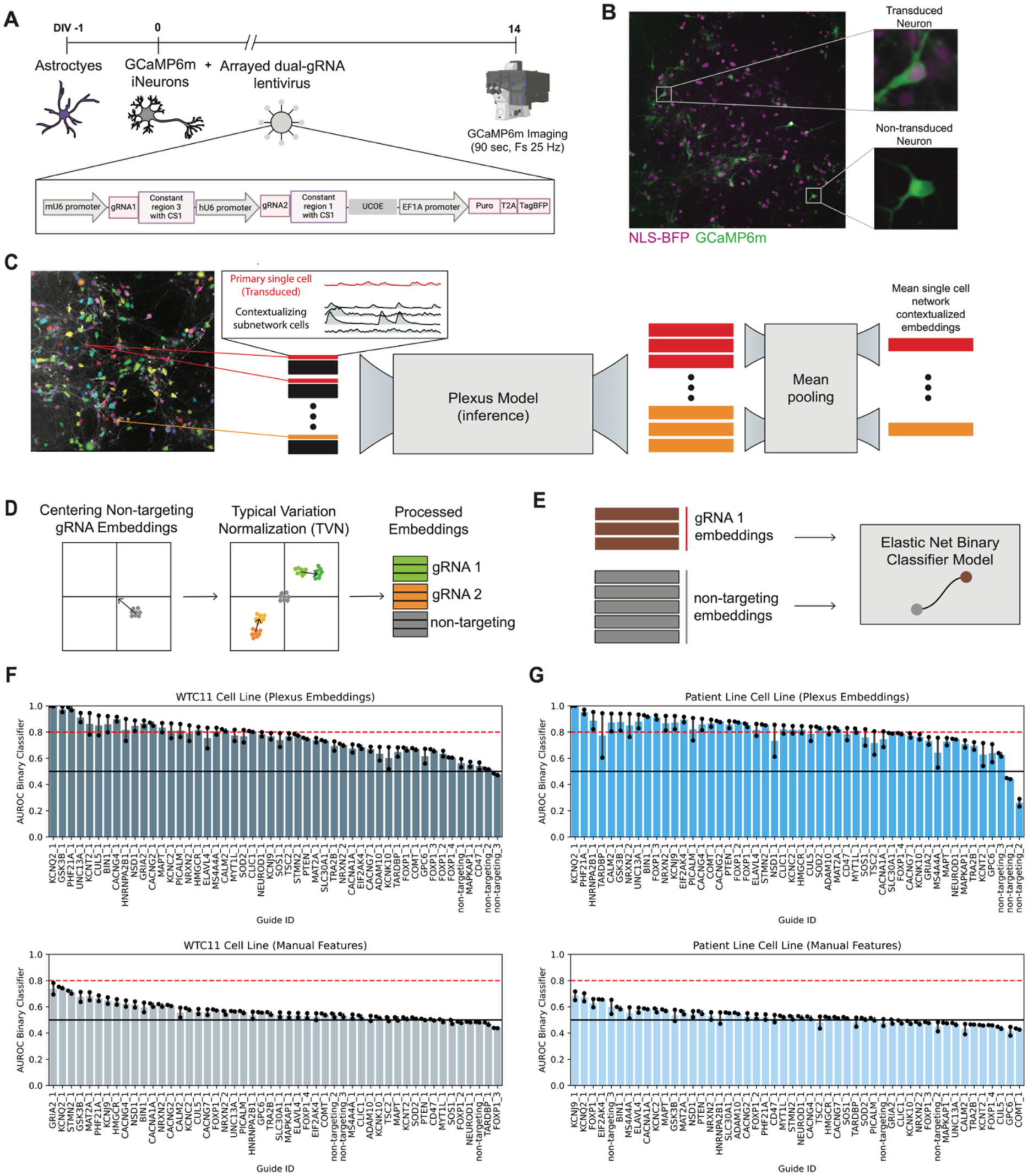
Arrayed CRISPRi screening using self-supervised neuronal activity dynamic phenotypic embeddings. (A) Illustration of the arrayed CRISPRi screening assay. (B) Example images of transduced and non-transduced GCaMP6m iNeurons. (C) Transduced cells have ten contextualizing subnetworks generated from the field-of-view video. These subnetworks are passed through the Plexus encoder to generate the network contextualized single-cell embeddings, which are then mean pooled to a single embedding per transduced cell. (D) Illustration of the embedding pre-processing technique of centering embeddings on the non-targeting control phenotypes and aligning across batches using typical variation normalization. (E) Illustration of elastic-net binary classifier task for the separability of non-targeting controls from gene knockdown phenotypes. (F) The area under the receiver operator curve for the elastic-net binary classification model for separating each dual guide vector vs non-targeting control trained on Plexus embeddings (top) and engineered features (bottom) for the WTC11 cell line (AUROCs are shown for both 4-well splits). The dotted red line represents the significant phenotype threshold of 0.8; the black line represents a random AUROC of 0.5. (G) The area under the receiver operator curve for the elastic-net binary classification model for separating each dual guide vector vs. non-targeting control trained on Plexus embeddings (top) and engineered features (bottom) for the Patient cell line.

After imaging all our neuronal cultures, we extracted the single-cell GCaMP signals from the videos and segmented the BFP+ nuclei using a fine-tuned CellPose^31^ model to ensure we were only investigating activity traces from transduced neurons (neurons that had a genetic perturbation). We then split the cells into transduced and non-transduced activity traces (**Methods**). We trained the Plexus model on these data for 1000 epochs. Plexus embeddings for these traces were generated by first filtering to only transduced neurons, generating ten subnetworks (sets) per transduced cell, and passing them through the Plexus model encoder. The embeddings from the ten contextualizing subnetworks or sets were then mean pooled per cell to generate network contextualized single cell embeddings (**Figure 4C**). The embeddings were then centered on the non-targeting controls and had cross-batch covariance structure aligned using typical variation normalization (**Figure 4D**) following the standard practice^32–34^ in phenotypic screening.

To determine genetic perturbation-driven phenotypes, we sought to find linear patterns in the embeddings that are generalizable across held-out wells (not driven by well-to-well stochasticity). To do this, we trained elastic-net binary classifier models to distinguish non-targeting control cells from those that had a genetic perturbation (**Figure 4E**). We performed a 50-50 train-test split with cross validation, such that all cells from the same well were in the same fold; this ensured the model did not overfit to well-specific information, such as synchronicity patterns or batch effects. A two-fold cross-validation was performed to ensure 4 wells per split. The AUROCs of the elastic-net models trained on the two-fold splits were used to measure phenotypic separability from non-targeting controls for the WTC11 (**Figure 4F**) and Patient (**Figure 4G**) cell lines, respectively. We found the Plexus embeddings captured 22.5 times (18X WTC11; 27X Patient) more significant phenotypes compared to signal processing-based features, as determined by the number of average AUROCs above 0.8. An AUROC of 0.8 was chosen as a threshold for meaningful phenotypes, as we believe this represents meaningful classification performance. This suggests that Plexus uncovers generalizable biological phenotypes that would not have been discovered using existing phenotyping methods. This represents an advancement in the field of neuronal activity phenotypic profiling, as it expands the number of gene perturbations that may be used to revert disease-related neuronal activity phenotypes.

### Validating CRISPRi Platform: Plexus identifies that *KCNQ2 knockdown drives a* distinct functional activity phenotype

To validate the ability of Plexus embeddings to capture biologically meaningful and interpretable phenotypes in our CRISPRi screening platform, we focused on the knockdown of the *KCNQ2* gene. This analysis acts as a positive control for Plexus and our CRISPRi screening platform. The loss of function of *KCNQ2* is known to drive dysregulated neuronal activity. Specifically, KCNQ2 encodes a voltage-gated potassium channel that helps regulate the M-current and whose loss of function has been implicated in childhood epilepsy ^35–37^.

To investigate distinguishing the *KCNQ2* knockdown-driven phenotype from non-targeting controls, we fit an elastic net regularized logistic regression ^38^model for each cell line. We performed a three-fold split on the single-cell embeddings from the non-targeting and *KCNQ2* knockdown cells. To demonstrate the performance of the elastic net models, we calculated the AUROC across the three-fold class-balanced splits (**Figure 5A**). To ensure that the model was not shortcut learning, we also performed a random shuffling of the labels and the embedding features and showed that in both cases, the elastic net AUROCs decreased to random chance (**Figure 5A**). We then took the mean of the elastic net model parameters from the three splits as a generalizable phenotype vector. Projecting the single-cell Plexus embeddings from all splits onto this mean predictive axis across the three-fold splits demonstrates phenotypic separability for both the WTC11 and the Patient cell lines (**Figure 5B**).

**Figure 5:**
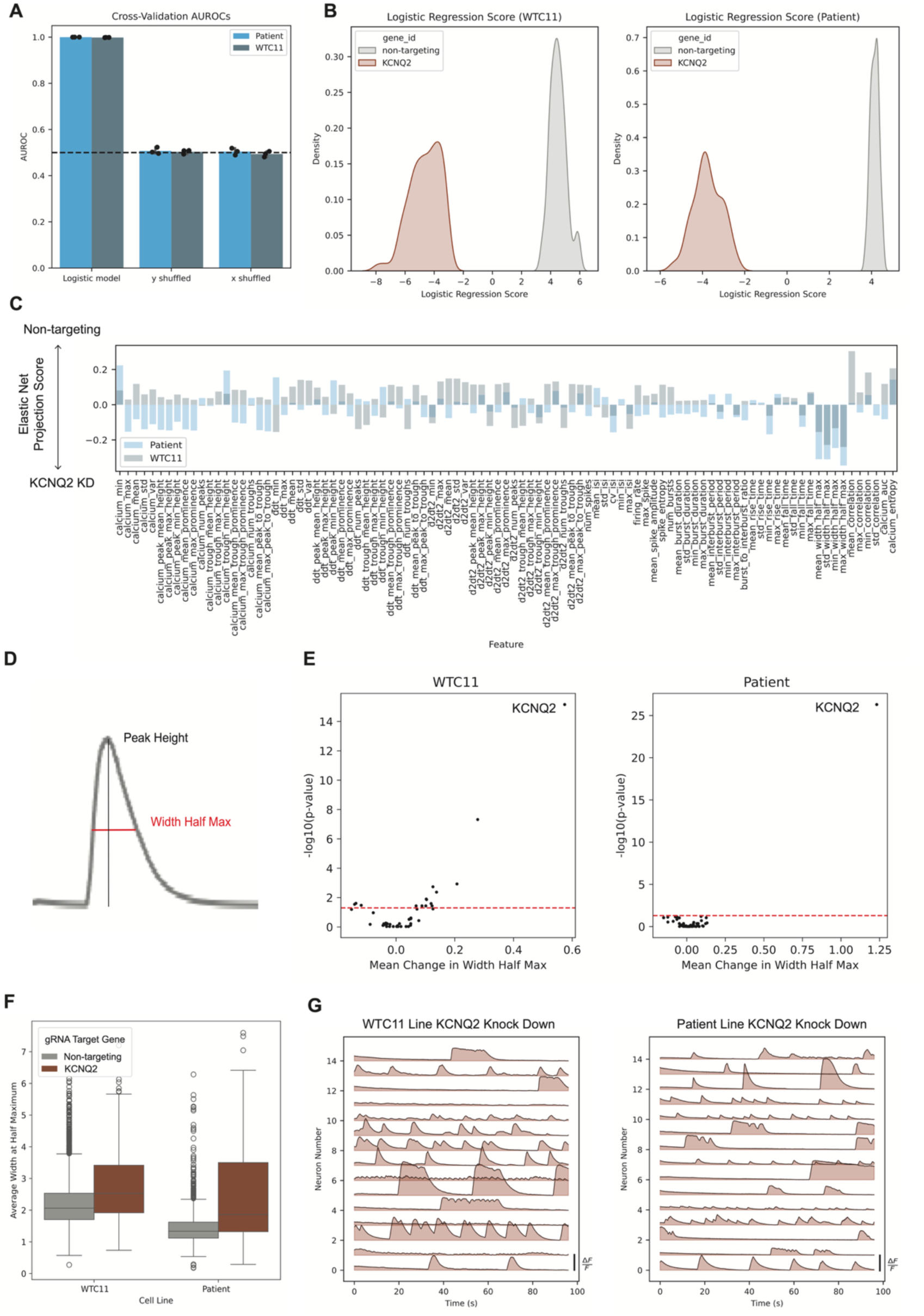
*KCNQ2* knockdown induces a distinct phenotype in two iPSC-derived iNeuron cell lines. (A) Elastic net model AUROC on three-fold cross-validation, showing the ability to separate *KCNQ2* and non-targeting control Plexus embeddings. Models with shuffled labels (y-shuffled) or shuffled Plexus features (x-shuffled) are not predictive, suggesting that no shortcut learning, batch-specific effects, or confounding feature correlations led to the separability of the phenotypes. (B) Projection of single-cell embeddings onto the mean elastic net projection axis of the OPLS models across three-fold splits reveals consistent separability between *KCNQ2* and non-targeting phenotypes in the WTC11 cell line (left) and the Patient cell line (right). (C) To interpret the Plexus embeddings, the mean predictive projection axes from both cell line models were projected onto the covariance matrix of Plexus embedding features and engineered features. The resulting predictive projection scores reflect feature influence: negative scores indicate a *KCNQ2* knockdown-like phenotype, while positive scores align with non-targeting controls. (D) Illustration of peak width at half maximum of a GCaMP6m calcium response peak. (E) Mean change in peak width at half maximum versus -log10 BH adjusted p-value from the Mann-Whitney U test, comparing the non-targeting control null distribution and gene knockdowns for the WTC11 cell line (left) and the Patient cell line (right). (F) Box plot (center line, median; box limits, upper and lower quartiles; whiskers, 1.5x interquartile range; points, outliers) showing the distribution of mean peak width at half maximum for both cell lines, with WTC11 (left) and Patient (right). The plot highlights differences in mean peak width half maximum between non-targeting controls and *KCNQ2* knockdown cells in each cell line. (G) Representative calcium response traces of cells with the most *KCNQ2* knockdown-like phenotypes, as identified by projection onto the predictive projection axis, show distinctive phenotypes for the WTC11 (left) and Patient cell lines (right).

Next, we performed an interpretability analysis to understand which manually engineered features were covarying with the Plexus features driving this KCNQ2 phenotype. In other words, we used the existing lexicon of engineered features to describe the patterns identified with the learned embeddings *post hoc*. To do this, we calculated the covariance matrix between the manually engineered features and Plexus embedding features across all cells in the screening dataset. We then took the mean predictive axis from the elastic net models and projected it onto the manually engineered features via the covariance matrix. This resulted in what we term the predictive projection score for each engineered feature (**Figure 5C**). Negative predictive projection scores correspond to engineered features whose increase in value is associated with the *KCNQ2* knockdown phenotype, while positive scores correspond to engineered features whose increase in value is associated with a non-targeting control phenotype. Notably, the Plexus features most associated with the KCNQ2 knockdown phenotype covary greatly with the peak width at half the maximum (PWHM) of calcium peaks (**Figure 5D**). We found that the importance of PWHM was consistent using other linear methods as well (**Supplementary Figure 5**). To assess the PWHM’s unique relationship with the *KCNQ2* knockdown phenotype, we performed Mann-Whitney U tests for each dual guide vector in the screen against the non-targeting null distribution with Benjamini-Hochberg (BH) multiple hypothesis correction. In both cell lines, this KCNQ2 knockdown phenotype is distinct from the non-targeting control and unique in this phenotype compared to the other genetic perturbations (**Figure 5E**). Although the PWHM is not sufficient to separate the non-targeting and *KCNQ2* knockdown alone, we do observe the increase in this engineered feature in both cell lines (**Figure 5F**). This distinct phenotype can be observed when visualizing cell traces that score the most *KCNQ2* knockdown-like when projected onto the predictive projection axis (**Figure 5G**).

### Exploring new biology: Uncovering genetic perturbations that map between disease phenotypes in a model of frontotemporal dementia

The goal of phenotypic screening is to define a ‘healthy’ and ‘disease’ phenotype and find perturbations that map cells between these states. To showcase our Plexus-enabled CRISPRi screening platform we focused on a genetic model of frontotemporal dementia. To this end, we used two isogenic cell line pairs generated using Cas9-mediated editing in the WTC11^39^ and Patient Line^30^ backgrounds resulting in pairs of cells with mutant heterozygous V337M MAPT (hereafter mutant tau) or Wild-type V337V MAPT (hereafter WT Tau) (**Figure 6A**). The Patient line was derived from a patient with the MAPT V337M mutation and symptomatic frontotemporal dementia and the isogenic line was generated by correcting the mutation to V337V^30^. The WTC11 line was derived from a healthy patient and was edited to obtain the MAPT V337M mutation^38^. This mutation in MAPT, encoding the protein Tau, was chosen as a disease model, as it is fully penetrant for frontotemporal dementia and has been shown in previous studies to drive aberrant neuronal activity in iPSC-derived neurons^38^.

**Figure 6:**
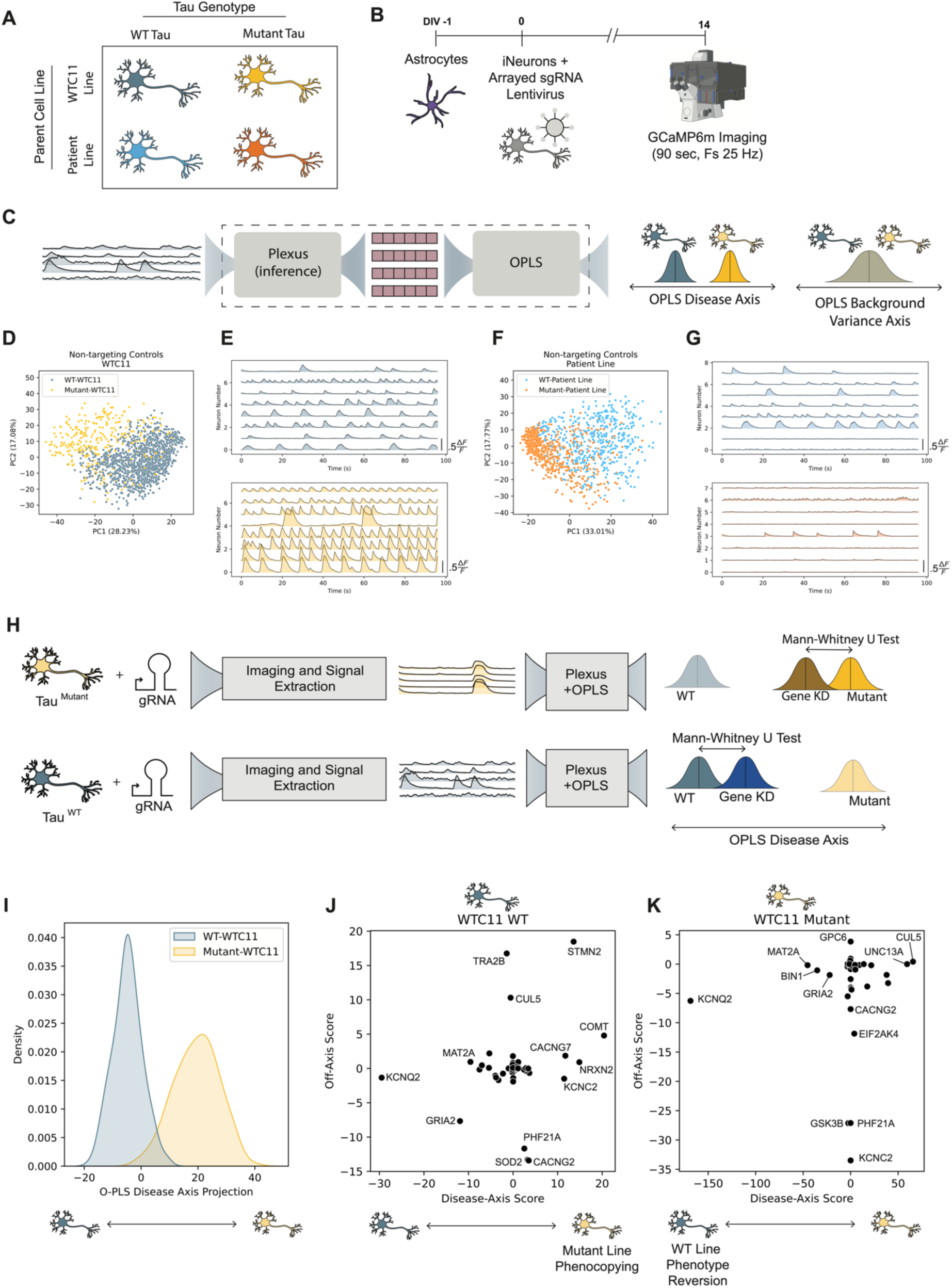
Uncovering gene knockdowns that map between distinct cell line phenotypes. (A) Illustration of the four cell lines used in arrayed CRISPRi screening. Two parent lines with distinct genetic backgrounds were used: WTC11 and Patient. Additionally, both cell lines had isogenic pairs with either the mutant V337M or WT V337V Tau genotype. (B) An illustration of the arrayed CRISPRi screening strategy. (C) To find genetic modifiers of activity that explain the variance between the WT and Mutant Tau neurons, we developed a pipeline comprised of the Plexus model in inference for embedding generation and an OPLS model for separating the embedding features into those that best explain the Tau genotype-associated phenotype (disease-axis) and the orthogonal cell-to-cell variation (off-axis). (D) The lower-dimensional projection of the first two principal components shows distinct activity phenotypes captured by the Plexus embeddings of the isogenic WTC11 cell line pairs transduced with non-targeting control dual guides. (E) GCaMP6m activity traces from a field-of-view video of the WT Tau (top) and Mutant Tau (bottom) WTC11 cell lines transduced with a non-targeting control dual-guide construct. (F) The lower-dimensional projection of the first two principal components shows distinct activity phenotypes captured by the Plexus embeddings of the isogenic Patient cell line pairs transduced with non-targeting control dual guides. (G) GCaMP6m activity traces from a field-of-view video of the WT Tau (top) and Mutant Tau (bottom) Patient isogenic cell lines transduced with a non-targeting control dual-guide construct. (H) The transduced Tau genotype-specific iNeuron activity traces associated with gene knockdowns are then run through the Plexus model to generate the network contextualized single-cell embeddings and projected onto the OPLS axis to score the phenotypic shift associated with gene knockdown. We then compare the distributions of the gene knockdown scores to the non-targeting control scores using a Mann-Whitney U test to determine statistically significant shifts along both the disease axis and the orthogonal off-axis. This provides insight into genes that shift the mutant Tau cell lines toward a WT phenotype and the WT Tau cell lines toward a mutant phenotype. (I) Projecting the non-targeting control single-cell Plexus embeddings onto the disease axis shows separability between the WT and mutant Tau WTC11 cell line phenotypes. (J) Plotting the disease-axis and off-axis scores suggests potential failure modes driving the mutant Tau phenotype. The scores represent the product of the -log10 BH-corrected p-values from the MWU tests and the shift in the barycenter between the gene knockdown distribution and the gene knockdown distribution for the WTC11 WT Tau cell line. Knockdowns that have low off-axis scores and positive on-axis scores represent potential genes that, when knocked down, play a role in the mutant Tau aberrant activity phenotype. (H) Plotting the disease-axis and off-axis scores for the WTC11 mutant line gives insight into potential genes that, when knocked down, could have therapeutic potential, specifically those with a low off-axis score and a large negative disease-axis score.

The CRISPRi screen targeting the 52 genes was performed in both pairs of isogenic cell lines, resulting in over 36000 single-cell neuronal activity traces with different mutant Tau statuses and genetic perturbations (**Figure 6B**).

To define our ‘disease’ and ‘healthy’ phenotypes, we leveraged our Plexus model. We trained the Plexus model for 1000 epochs on the data from this CRISPRi screen and generated single cell embeddings as previously explained above. To demonstrate the clear distinction in the activity phenotypes, we visualized the two - dimensional principal component projection of the Plexus embeddings and plotted representative activity traces (WTC11: **Figure 6D-E**; Patient: **Figure 6F-G**).

After determining the phenotypic states that we wished to map between, we sought to determine genes that when perturbed map between the two states with little off-target effect. To deconstruct the off-target and on-target axes of phenotypic change, we fit an OPLS^37^ model on the Plexus embeddings from cells with non-targeting control guides, with Tau status as the response variable. An OPLS model was fit for each isogenic cell line pair using a 50-50 train-test well-aware split. This resulted in a disease axis (predictive axis) describing the WT and Mutant Tau differences in activity, and an off-target axis (orthogonal axis) describing all orthogonal variation in activity (**Figure 6C**; **Methods**). The projection of the test split non-targeting control Plexus embeddings onto the OPLS disease axis shows generalizable separability between the WTC11 mutant tau and WT tau line phenotypes (**Figure 6I**)

To determine gene perturbations that shift cells from a mutant Tau phenotype towards a WT tau phenotype, we projected the Plexus embeddings corresponding to all perturbed mutant Tau cells onto the one-dimensional disease and off-target axes. We then compared the disease axis and off-target axis distributions between the non-targeting control mutant Tau cells and the cells with each gene perturbation using Mann-Whitney U tests (**Figure 6H**). We use the product of the mean shift along the disease or off-target axis and the -log10 BH corrected p-values to calculate the on-target disease score and off-target scores. By plotting the disease-axis score versus the off-axis score were able to determine genes that, when perturbed, have a high on-target effect and a low off-target effect (**Figure 6J-K**). These correspond to either potential failure modes in disease that, in a WT background, drive an aberrant activity phenotype or potential therapeutic targets that, when knocked down in the mutant line, map back to a healthy phenotype.

In the case of the mutant Tau cell line, we see that *MAT2A*, *BIN1*, and *GRIA2* are targets that, when knocked down, shift the phenotype closer to a healthy phenotype while minimizing off-target activity (**Figure 6K**). Although *KCNQ2* has a higher on-axis score shift, it also corresponds to a larger off-axis score driven by the phenotype observed in **Figure 3**. Although neither the knockdown of *MAT2A*, *BIN1*, nor *GRIA2* completely rescued the mutant Tau phenotype, this framework provides insight into genes and their related pathways that may play a role in driving the aberrant neuronal activity phenotype. Traditional signal processing features showed limited phenotype detection capabilities. While detecting *BIN1* and *GRIA2* with reduced effect sizes, this approach missed *MAT2A* and failed to identify *KCNQ2*’s known off-target effects—a critical oversight given *KCNQ2*’s established role in disease-relevant phenotypes (**Supplementary Figure 7**). These limitations highlight the superior detection capability of OPLS models leveraging Plexus embeddings.

## Discussion

We introduced an integrated experimental and computational phenotyping method for high-content screening of neuronal activity dynamics. We demonstrated that our Plexus self-supervised learning model outperforms existing methods at capturing and differentiating neuronal activity phenotypes for screening. Plexus is particularly performant at distinguishing small changes to underlying dynamics, which we demonstrated via *in silico* perturbations to network dynamics simulated using a multivariate self-regulating Hawkes process. This performance was further demonstrated on experimental data of our iPSC-derived neuronal culture, where Plexus outperformed existing methods at determining phenotypes driven by neuroactive compounds. Applying Plexus to CRISPRi screening demonstrated the discovery of a greater number of generalizable biological phenotypes driven by gene knockdowns.

Extending this approach to a MAPT mutant model of aberrant neuronal activity, we showed the ability to use the learned embeddings from Plexus to determine potential disease-specific modifiers of neuronal activity dynamics.

Plexus opens the possibility of large-scale phenotypic screening for genetic modifiers of neuronal activity in homeostasis and disease. Additionally, while we focused on a model of neurodegeneration in this study, this approach can be extended to model any disease with an aberrant neuronal activity phenotype. Combining genetic and small molecule perturbations with Plexus phenotyping could also provide insight into the mechanism of action of neuroactive compounds – a challenging task in drug discovery efforts. While we focused on neuronal activity as a phenotype, this method could be applied to any dynamic biological process where biological context influences single-cell phenotypes.

Our experimental iPSC-derived iNeuron and astrocyte co-culture model enabled us to study complex cellular dynamics *in vitro*. While our model focused on excitatory cortical-like neurons, extending this system to include inhibitory interneurons could provide a useful system for studying excitatory/inhibitory imbalance, which is implicated in many neurological diseases. Additionally, with recent advancements in modeling region-specific neuronal subtypes from iPSCs, Plexus could be used to extend genetic modifier screens to uncover genes with subtype-specific regulatory effects. For example, iPSC-derived motor neurons could be used to determine genes involved in the aberrant activity phenotypes associated with ALS. More broadly, our study highlights the potential of combining targeted experimental platforms with advanced machine learning techniques to generate therapeutic hypotheses for complex diseases.

Several technical considerations and limitations should be noted. While calcium imaging with GCaMP6m provides a scalable readout of neuronal activity, it captures only a subset of neuronal function via the secondary messenger of calcium. Future studies combining our approach with electrophysiological recordings or voltage sensors could provide additional validation and mechanistic insights. Additionally, while our arrayed screening approach enables precise phenotypic measurements, it is limited in throughput compared to pooled screening methods. The trade-off between depth and breadth of phenotypic characterization remains an important consideration for future screening efforts. And finally, the current experimental system relies fully on spontaneous activity. This presents the opportunity to integrate optogenetic tools capable of stimulating neuronal activity, which could provide valuable insight into disease mechanisms that only arise under high levels of activity.

Overall, we believe this study demonstrates the promise of leveraging biology-aware SSL methods to study complex phenotypes and their genetic regulation at scale. We acknowledge that this work stands on the shoulders of incredible efforts in leveraging self-supervised learning for phenotyping static cellular states, such as transcriptomics and morphology. We hope that our study will spur further development of systems that combine iPSC-derived models of cellular processes, perturbation biology, and SSL to further large-scale phenotypic screening efforts.

## Methods and Materials

### Human iPSC culture (adapted from Tian et al. 2021)

The WTC11 WT and heterozygous MAPT V337M iPSC lines were gifts from Dr. Li Gan’s Lab. The GIH6C1 WT and heterozygous MAPT V337M iPSC lines were acquired from the Tau Consortium Stem Cell Group. Human iPSCs from the WTC11 background (Male) and the GIH6C1 background (Female)^30,40^ (referred to as the Patient line in the main text) were cultured in StemFlex Medium (GIBCO/Thermo Fisher Scientific; A3349401). Cell lines containing the iPSCs were grown in tissue culture grade plates or dishes coated with Growth Factor Reduced, Phenol Red-Free, LDEV-Free Matrigel Basement Membrane Matrix (Corning; 356231) diluted 1:100 in Knockout DMEM (GIBCO/Thermo Fisher Scientific; 10829-018). StemFlex Medium was replaced daily.

When cells reached 80% confluency, cells were dissociated with StemPro Accutase Cell Dissociation Reagent (GIBCO/Thermo Fisher Scientific; A11105-01) at 37oC for 5-7 mins, centrifuged at 250 xg for 5 min, resuspended in StemFlex Medium supplemented with 10 nM Y-27632 dihydrochloride ROCK inhibitor (Tocris; 125410) and placed onto Matrigel-coated plates or dishes. hiPSCs were maintained in Y-27632 until colonies reached greater than 15 cells per colony. Studies at UCSF with human iPSCs were approved by the Human Gamete, Embryo, and Stem Cell Research (GESCR) Committee. Informed consent was obtained from the human subjects when the hiPSC lines were originally derived.

### GCaMP6m cell line engineering

To generate the GCaMP6m iNeurons, the hiPSCs for all cell lines were transduced with a lentivirus delivering the GCaMP6m construct (gift from Dr. Michael Ward). The cells were transduced at an MOI of 5, followed by FACs sorting for the top 25% highest expressing cells, resulting in polyclonal populations of GCaMP6m+ NGN2 hiPSCs for each unique cell line (WTC11 and Patient cell lines with WT and Tau mutation statuses).

### Arrayed dual-guide lentivirus generation

Arrayed lentivirus was generated by an optimized reverse transfection protocol in 96-well plate (Corning, #3628). Each arrayed plasmid was constructed using the *NEBuilder® HiFi DNA Assembly Master Mix (NEB; E2621L) and NEB 5-alpha Competent E. coli (NEB; C2987P)* and purified with the *QIAprep spin miniprep kit (Qiagen; 27106*). An envelope plasmid (MD2G; Addgene 12259) and a manipulated packaging plasmid (psPAX2-long-pCMV) were purified with the ZymoPURE II Maxiprep kit (Zymo Research; D4203). All plasmids were quantified with the Nanodrop.

After dispensing 22.5 μl of a master mix containing 20 μl of the Opti-MEM (Thermo Fisher scientific; #31985088), 100 ng of packaging plasmid, and 10 ng of envelope plasmid on each well, 1.25 μl of arrayed plasmids (100 ng/ul) were added first. 12.95 ul of transfection master mix composed of 12.24 μl of 1 mM HEPES (pH 7.3, Thermo Fisher Scientific; 15630130) and 0.71 μl of TransIT-VirusGENE (Mirus; MIR 6700) was added into each well, vortexed at 1,500rpm at room temperature for 15 seconds using the BioShake IQ (Bulldog Bio; 1808-0506) immediately. After incubation for 35 min, 100 μl (9e4 cells) of trypsinized Lenti-X cells (TakaraBio; 632180) was dispensed into each well, mixed well at 1,500 rpm for 15 seconds using the BioShake IQ again, and incubated at room temperature for 1 hour before transferring to a tissue culture incubator (37 °C, 5% CO2). 15.5 hours post transfection, the spent media was removed carefully and 119 μl of a serum-free Pro293a-CDM media (Lonza; BEBP12-764Q) with the ViralBoost (AlstemBio; VB100) was added. Lentivirus was harvested 43 hours post reverse transfection, and aliquots were stored at −80 °C.

### Dual-guide RNA cloning

Guide sequences targeting the transcription start sites of the 52 target genes were synthesized by IDT in an arrayed formate Dual guide RNAs were cloned in a lentiviral vector backbone using Gibson assembly cloning. An insert DNA sequence with flanking *BsmBI* restriction sites and containing a constant region with Perturb-seq compatible capture sequence 1 (CS1: GCTTTAAGGCCGGTCCTAGCAA) was cloned into an intermediate transfer vector and transformed into bacteria for plasmid amplification (See insert sequence below). To prepare a linear insert for cloning, the amplified plasmid was digested using *BsmBI* restriction digestion, and the desired fragment was purified using gel purification. The backbone vector was also linearized using *BsmBI* restriction digestion. This backbone vector (pLGR134) was derived from pLGR002 (Addgene #188320) by swapping the *BstXI* and *BlpI* sites for a pair of *BsmBI* sites. ssDNA oligos for guides were ordered with 20 base pair overhangs on both 5’ and 3’ sides, which partially overlap the constant region and mU6 promoter for guide 1 and constant region and hU6 promoter for guide 2. Linearized vector, insert, and guide oligos were then assembled into dual guide vectors in a Gibson assembly reaction. Colonies were validated by Sanger sequencing.

Insert sequence before BsmBI restriction digest: CGTCTCAagaggtttcAGAGCTAAGCACAAGAGTGCATAGCAAGTTGAAATAAGG CTAGTCCGTTTACAACTTGGCCGCTTTAAGGCCGGTCCTAGCAAGGCCAAGT GGCACCCGAGTCGGGTGCTTTTTTTGCTCGAATCTACACTCAGCTATGGCGC GCCCCAAGGTCGGGCAGGAAGAGGGCCTATTTCCCATGATTCCTTCATATTTG CATATACGATACAAGGCTGTTAGAGAGATAATTGGAATTAATTTGACTGTAAAC ACAAAGATATTAGTACAAAATACGTGACGTAGAAAGTAATAATTTCTTGGGTAG TTTGCAGTTTTAAAATTATGTTTTAAAATGGACTATCATATGCTTACCGTAACTT GAAAGTATTTCGATTTCTTGGCTTTATATATCTTGTGGAAAGCCAgaaacatgGAA AGGAGACG

### Human organoid-derived astrocyte generation

Control iPSCs from a healthy donor (WT#2: 36yo Asian Female, ATCC-KYOU-DXR0109B) were differentiated into spherical neurospheres (SNMs) containing neuroectodermal progenitors which were differentiated toward an astrocytic lineage following a previously published protocol (Serio et al., 2013). First, iPSC cultures were cultured with StemFlex (Gibco) to reach 80% confluence. On Day 0, iPSCs were dissociated to single cells with Accutase (Inno Cell Tech) and 2 million cells were transferred to one 125mL Spinner bottles (Corning, CLS3152) on a Duramag Spinner (Chemglass, CLS-4100-09, 65rpm) with Neural Induction media (DMEM/F12 (Gibco), 1% N2 (Thermo), 0.1% B27 (Gibco), 1% NEAA (Thermo), 1% Glutamax (Gibco) supplemented with SB431542 (R&D Systems), LDN193189 (StemGent) and Y-27632 (, R&D Systems). After 24 hours of spinning in suspension, the cells formed Embryoid Bodies (EBs), where half of the media was changed to fresh Neural Induction media, removing half of the Y-27632. The next day, a full media change was performed. On day 4, the media was changed to Rosette Formation Media (half DMEM/F12 half Neurobasal, 0.1% B27, 1% NEAA, 1% Glutamax, supplemented with SB431542, LDN193189 and 20 ng/mL FGF-2 (R&D Systems). After day 10, the media was changed to Neural Progenitor Media (Neurobasal, 0.1% B27, 1% NEAA, 1% Glutamax, supplemented with 20 ng/mL FGF-2). After day 26, the EBs were grown in suspension for another 25 days with Induction Medium (Neurobasal, 0.1% B27, 1% NEAA, 1% Glutamax supplemented with 20 ng/mL LIF (Peprotech) and 20 ng/mL EGF (R&D Systems), and then for further 20 days with Propagation Medium (Neurobasal, 0.1% B27, 1% NEAA, 1% Glutamax containing 20 ng/mL FGF-2 and 20 ng/mL EGF) becoming SNMs. To harvest the astrocytes into single cells, SNMs were incubated with Papain (Worthington) for 15 minutes at 37°C and were mechanically dissociated using a p100 pipette and plated on Geltrex-coated plates as a monolayer. The monolayer of neural progenitors was cultured for 14 more days in Propagation Medium and then for another 14 more days in Astrocyte CNTF medium (Neurobasal, 1% Glutamax, 1% NEAA, 0.2% B27 supplement, 10 ng/mL CNTF (R&D Systems), at which the cells were considered astrocyte progenitors (day 100). The astrocyte progenitors were successfully frozen in a Freezing Medium (AMSBIO Stem-CellBanker) and stored in liquid nitrogen. Part of the culture was passaged another four times to increase maturity and sorted for the mature Astrocytes marker CD49f (BioLegend; 313615; used at 1:200 dilution) on the Sony Cell Sorter (Sony; SH800S) to filter out any invasive cells. Pure astrocyte cultures were kept in culture for more than 200 days to reach mature astrocyte identity and then cryobanked in 10% DMSO for later use.

### iNeuron differentiation and co-culture with hiPSC-derived astrocytes

hiPSCs were pre-differentiated into day 0 iNeurons, according to Tian et al.^20^, and frozen down with 10% DMSO for cell banking. At day-1 of the iNeuron differentiation hiPSC-derived astrocytes were plated onto 384-well plates (Grenier; 142761) in BrainPhys neuronal media (composed according to Bardy et al. 2015)^41^ at 6k cells per well with Growth Factor Reduced, Phenol Red-Free, LDEV-Free Matrigel Basement Membrane Matrix (Corning; 356231) diluted 1:200 in Knockout DMEM (GIBCO/Thermo Fisher Scientific; 10829-018) using the BRAVO liquid-handling robotic system (Agilent). For CRISPRi screening, iNeurons were transduced at day 0 via reverse transduction, where the lentiviral aliquots were stamped into randomized locations in the 384-well plates using the BRAVO liquid-handling robotic system (Agilent) prior to the addition of D0 neurons. D0 neuron vials were removed from liquid nitrogen and defrosted at 37°C for 2 minutes and resuspended in BrainPhys neuronal media (composed according to Bardy et al. 2015)^41^ with 2 μg/ml doxycycline (Takara; 631311). The iNeurons were then spun down at 300 xg for 5 minutes, at which point the spent media was removed, and the cells were resuspended in fresh BrainPhys neuronal media with 2 μg/ml doxycycline. The cells were then plated onto the 384-well plate using the BRAVO liquid-handling robotic system. Three days after plating, half of the culture media was removed, and fresh BrainPhys media with 6 μg/ml 5-Fluoro-2′-deoxyuridine (5FdU) (Millipore Sigma; F0503) and no doxycycline was added. After day 3, a half-change for supplementation with fresh BrainPhys media with no 5FdU or doxycycline was performed twice weekly using the BRAVO liquid handling robotic system.

### GCaMP6m imaging

Calcium activity was recorded on day 14 and day 21 using a wide-field Eclipse Ti2 microscope (Nikon) and the NIS Elements software version 5.30.05. A 20x water immersion objective, 4×4 16-bit binning, an exposure of 40 ms, or a sampling frequency of 25 Hz was used during imaging. An ORCA-Fusion BT Digital CMOS camera (Hamamatsu; C15440-20UP) was used for photon detection and a spectra light engine (Lumencore) was used for the epifluorescence light source. In each well of the 384-well plates 2 videos were recorded from two fields of view separated in x by 0.9 mm and vertically centered in the well. Each video was recorded for 96 seconds, corresponding to 2400 individual frames. In addition to each GCaMP6m video, an image of the nuclear-localized BFP with the DAPI channel filter was taken during the CRISPRi screening for downstream use in determining single-cell transduction.

### Neuroactive stimulation of neuron astrocyte co-cultures

To elicit distinct neuronal activity phenotypes, we treated D13 co-cultures with 1.5 μM TTX (Tocris, 1078) or 1 nM TeNT (Sigma, T3194) and incubated for 24 hours before imaging. For stimulation with 2 mM Ca2+ and 2 mM Mg2+ we supplemented BrainPhys media with either additional CaCl2 (Millapore Sigma; C4901) or additional MgSO4 (Millipore Sigma; M7506) to increase the concentrations of Ca2+ from 1.1 to 2 mM and Mg2+ from 1 to 2 mM respectively. A complete media change to the 2 mM solutions was performed 1 hour before imaging to elicit modified activity without impact on neuronal survival.

### Immunostaining and Imaging Astrocyte iNeuron co-cultures

iNeurons and Astrocytes were co-cultured according to the description above until DIV14, when the cultures were fixed with 4% paraformaldehyde (diluted from a 16% solution with DPBS; Electron Microscopy Sciences, 15710) for 15 minutes at room temperature. After washing three times with DPBS, blocking and permeabilization were performed with DPBS + 3% BSA + 0.1% Triton X-100 (Millipore Sigma, X100) for 30 minutes at room temperature. Primary antibodies against GFAP (1:500, rabbit polyclonal; Thermo Fisher Scientific, PA1-10019) and S100β (1:500, mouse monoclonal; Millipore Sigma, S2532), were then added in blocking buffer and incubated overnight at 4°C. Afterward, the samples were washed with DPBS + 0.1% Triton X-100 three times, incubated with pre-adsorbed secondary antibodies (1:500 goat anti-mouse IgG AF647 and 1:500 goat anti-rabbit IgG AF555; Abcam ab150119 and ab150086) for 1 hour at room temperature, washed two times with DPBS + 0.1% Triton X-100, incubated with 1 μg/ml of Hoechst (Thermo Fisher Scientific, H3570) for 20 minutes at room temperature and then washed two times before imaging. Samples were imaged with an ImageXpress Confocal HT.ai (Molecular Devices) microscope, using an Apo LWD 20x/0.95NA WI objective, 1×1 binning, and 100–400-ms exposure.

### Neuronal activity spiking activity phenotype simulation

The activity of a neuronal network was simulated using a self-regulating multivariate Hawkes process with discrete-time dynamics that incorporated excitation, negative feedback, and network connectivity. The network topology was generated using a Watts-Strogatz graph model, which allowed for adjustable clustering and rewiring probabilities to represent the small-world properties often observed in biological neural systems. The connectivity between nodes was captured in the adjacency matrix of the graph, with self-loops added to account for self-excitation effects. The simulation was conducted over a discrete time grid, where the total simulation time of 288 seconds was divided into bins of 0.04 seconds to match the sampling rate of the GCaMP imaging assay. The intensity function for each node evolved as a combination of baseline activity, excitation from neighboring nodes, and self-regulating negative feedback.

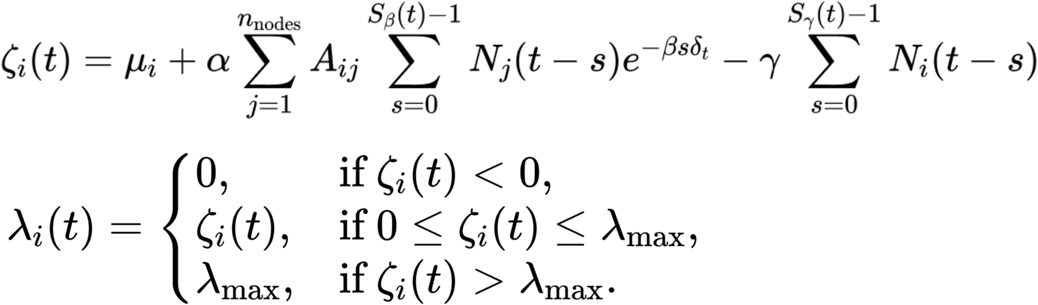

Baseline intensities μ_i_ were drawn from a normal distribution with a specified mean and standard deviation, introducing variability in the intrinsic firing rates of individual nodes. Excitation was modeled using an exponential decay kernel. At each time step, the recent spiking activity of the kernel was scaled by the synaptic strength parameter α and summed to compute the contribution of excitation to the intensity. Negative feedback was incorporated by linearly reducing the intensity based on the node’s own recent activity in the past *S*_β_ time steps. The strength and duration of this feedback were controlled by the parameters γ and *S*_γ_, respectively. The intensity at each time step was constrained to remain within a specified range, ensuring non-negativity and limiting the maximum number of events to roughly 250 events per second. The ∼250 events per second limit was calculated by computing the cumulative density of a Poisson distribution with a rate λ*_max_* of 0.4 δ_t_. Spiking event numbers for each node or cell were generated at each time step by sampling from a Poisson distribution with a rate λ(*t*) ∗ δ_t_. This procedure iteratively updated the event history and intensities for all nodes across all time steps, resulting in an array of spike events of the shape number of cells by total discrete time bins. Using this method, eight distinct neuronal activity phenotypes were then generated for 288 seconds. To match the neuronal activity data generated from the *in vitro* assay, we randomly sampled 96 seconds of the full simulation between 96 and 288 seconds. This was done to allow the model to reach steady-state behavior and induce stochasticity into the phase of the event timing. All the code for this simulation framework is available on GitHub in the plexus-simulate repository.

### Computational time-series extraction from GCaMP6m videos

To extract single-cell time series arrays from a single field-of-view video, the video first had a summed intensity projection (SIP) calculated along the time axis of the video. The SIP image was then normalized by subtracting the minimum and dividing by the image’s maximum to be scaled between 0 and 1. The array was then transformed pixel-wise dividing it by the exponentiated normalized SIP image to reduce the contrast in the image. A fine-tuned CellPose^31^ model was used to segment the neuronal soma from each field-of-view video, creating a single-cell mask with each identified cell having a unique number. This single-cell mask was then expanded to a tensor, each cell in its own channel. This single-cell tensor then had the x and y dimensions flattened, and the dot product of the tensor with the flattened video was performed to extract the single-cell time series. These time series were then normalized by the total number of pixels per single-cell mask to generate what we refer to as the raw time series. To generate the *ΔF/F* traces used for downstream analysis, each single cell trace was normalized by dividing by its baseline. The baseline was calculated using an asymmetric least-squares regression for each single cell trace with a smoothing parameter of 1e5 and an asymmetry parameter of 0.001. Inferred spike traces were then calculated using the OASIS^45^ deconvolve function with a sparsity penalty of 2, which were used for the engineered features. This process was then performed for all the field-of-view images for each 384-well plate, with the SIP image, the raw traces, the *ΔF/F* traces and the inferred spiking all saved to a hierarchical Zarr file for downstream processing and use.

All the code for this simulation framework is available on GitHub in the Plexus-extract repository.

### CellPose Model Fine-Tuning

Two CellPose models were fine-tuned for (1) neuronal soma segmentation and (2) nuclei segmentation. Cell segmentation masks were hand-annotated for 30 GCaMP6m summed intensity projection images to fine-tune the neuronal soma segmentation model. The soma model was fine-tuned starting with the weights of the pre-trained ‘cyto’ model and trained for 200 epochs using the default CellPose loss function. To train the model to segment the nuclear-localized BFP signal, 30 images were hand-annotated as training data. The nuclei model was fine-tuned starting with the weights of the pre-trained ‘nuclei’ model and trained for 200 epochs using the default CellPose loss function. The Cellpose Model weights are available in the Plexus-extract repository.

### Filtering neuronal activity data for transduced neurons

For the CRISPRi screening, each field of view video had a corresponding image of the DAPI channel to quantify which cells were NLS-BFP+, suggesting the cell had received the viral vector encoding for the guide RNA. When the NLS-BFP or nuclei images were present with the GCaMP6m videos, the nuclei images were segmented using a second nuclei-specific fine-tuned CellPose model. Neurons were then determined to be transduced if a nucleus mask greater than 75 was determined to be within the neuron soma segmentation mask. This resulted in a Boolean array corresponding to which cells were transduced or not per field of view, which was also saved to the hierarchical Zarr file generated during single-cell time-series trace extraction.

### Calculating the manually engineered multiparametric features

To generate multiparametric engineered features from the *ΔF/F* traces or signals, we started by estimating the first and second derivatives with respect to the time of the signal using numpy’s gradient method. For each cell, features were extracted from the single-cell signals and their derivatives to capture statistical, temporal, and synchrony characteristics. Basic signal statistics, such as minimum, maximum, mean, and variance, were computed alongside metrics for peak and trough properties, including peak height, prominence, and counts. Spiking activity was analyzed for inter-spike intervals (ISI) to assess firing regularity, and bursts were identified as groups of closely spaced spikes. Calcium signal dynamics, including rise times, fall times, and half-widths of peaks, were derived to understand the temporal behavior of calcium transients.

Network-level synchrony was quantified through correlations across cells, capturing interactions within the cell population. Additionally, the entropy of the calcium signal was calculated to assess variability, while the area under the curve (AUC) represented cumulative fluorescence. This extracted feature set was organized into a comprehensive data frame, which was used for downstream analysis. The code used to extract the multiparametric features is available in the Plexus GitHub repository.

### Training the Network-Aware Masked Autoencoders on simulated datasets

To train the Plexus, we started by collating the Zarr file containing the simulated GCaMP6m dataset and generating a Pytorch Dataset for the Plexus. To test the effect of different sized sets (subnetworks), we created datasets that generated random subnetworks of either 1, 4, 8, or 16 cells per simulated field of view. To generate the inputs for tokens we generated time chunks of 24-length windows corresponding to 0.96 seconds of the activity trace. These time windows were projected to 768 dimensions using a learnable linear transform, resulting in 100 tokens per cell. Each of the cells in the subnetwork had a learnable ‘position embedding’ corresponding to the ordinality of the time windows in every single cell. Then, each cell’s 100 tokens had a permutation invariant cell-specific learnable embedding added to allow the model to capture cell identity. The permutation invariant cell embeddings were initialized as random learnable vectors at the beginning of training. In each forward pass through the model, the unmasked tokens of each cell were mean-pooled into a single mean token per cell. The rank order of the L2 norm of these mean tokens was then used to assign each cell to one of the learnable vectors; the vector with the highest L2 norm was always assigned to the first learnable vector, and the mean token with the lowest L2 norm was always assigned to the last. This ensured that each cell, regardless of its position in the subnetwork, always received the same cell embedding. Then, the tokens corresponding to all the cells in the subnetwork were concatenated to make network-level inputs into the model. One CLS token and five register tokens were concatenated to the network-level input. We then masked 50% of the tokens and fed the unmasked tokens through the encoder to generate the latent unmasked embeddings. The masked tokens were added back via concatenation to the latent unmasked embeddings and fed through the model decoder. The decoded masked tokens were then transformed into the input time space via a learnable linear layer and compared to the time windows to calculate the mean squared error reconstruction loss. The variations of the model were each trained for 1000 epochs with a linear warmup of 15 epochs with a starting learning rate at epoch zero of 1e-6, a maximum learning rate of 5e-5, followed by a cosine decay to 1e-6. We chose 1000 epochs, as we found that it was sufficient for the model to converge (Supplementary Figure 1). The model was trained with a well-aware 80-20 train-validation split to ensure that the model was not learning well-specific bias. Each variation of the model was trained on a single NVIDIA TITAN RTXs with 24 GB for between 4 and 24 GPU hours. All the code used for training the Plexus models is available in the Plexus GitHub repository.

### Training the Network-Aware Masked Autoencoder on neuroactive simulation and CRISPRi screening data

To train the Plexus on the CRISPRi and neuroactive treatment datasets, we started by collating the Zarr files containing the 384-well plate with the cell treatments. To train the model, we randomly sampled subnetworks (sets) of 8 cells from the same field of video and generated time chunks of 24-length windows corresponding to 0.96 seconds of the activity trace. (8 cells were chosen for subnetworks, as this maximized the number of cells that could be trained across all four cell lines used in the CRISPRi screen. The Mutant Tau WTC11 line had fewer cells per field of view than the WT Tau WTC11, but on average, had at least 8 cells per field of view. If more cells per subnetwork were wanted, the seeding density of the Mutant Tau WTC11 cells would need to be increased.) This resulted in 50 chunks or time windows per cell. The model was trained following the same methods for the simulated data described above. However, this model was trained across 4 NVIDIA TITAN RTXs with 24 GB of memory each for 78 GPU hours.

### Generating the single-cell network contextualized embeddings

To generate the single-cell network contextualized embeddings, each single cell in each field of view of the image had 10 random subnetworks (sets) of 7 other cells sampled, such that the same 8-cell subnetwork was not sampled twice. This resulted in subnetworks with the cell of interest and 7 contextualizing cells from the same field of view. These 8-cell subnetworks were then passed through the encoder of the Plexus pre-trained model with no masking. In the case of the CRISPRi screen, only transduced cells were treated as cells of interest. To create the single cell embedding, the tokens from the cell of interest were averaged to create a single embedding dimension of 768. The 10 single-cell embeddings corresponding to each cell of interest were then averaged to get a single embedding per cell. All the code used for model inference can be found in the Plexus GitHub repository.

### Linear probing for classification of simulated neuronal activity phenotypes

Linear probing was performed on embeddings generated from the 1-cell MAE model, the 4-cell, 8-cell, and 16-cell Plexus models, and the manual features with and without the network-related manual features. Multinomial logistic regression was used for linear probing, solved using the lbfgs solver. The linear probing was performed on a 3-fold cross-validation split, and accuracy, as well as class-specific F1 scores, were calculated.

### Training and evaluating elastic-net classifiers for neuroactive treatment

These elastic-net classifiers were trained on the Plexus embeddings and the engineered features. Before training the classifiers, a three-way cross-validation split was performed in a well-aware manner, such that all the cells from the same well were in the same split. For each cross-validation train-test split, the data were scaled per feature to unit variance and zero mean on the train split. The elastic net binary classifier models were trained with equal weight on the L2 and L1 norm hyperparameters. To ensure class balance, the class with the most cells was randomly sampled to have the number of cells corresponding to the minority class. This resulted in meaningful AUROC curves without undue influence from class imbalance. The area under the receiver operator curves was calculated for each cross-validation split, and the mean ROC was plotted along with the standard deviation over the split.

### Preprocessing the gene knockdown Plexus embeddings

When the single-cell GCaMP6m time series traces were extracted, they were extracted into 8 Hierarchical Zarr files corresponding to each 384-well plate imaged. For each cell, the Plexus single cell embedding was generated as described above. The non-targeting control embeddings were then collated, and all the embeddings per plate were centered and scaled to the non-targeting control cells. Then, typical variation normalization was employed to align the mean and covariance structure of the non-targeting control embeddings across batches. Each batch was considered a plate in the screen, and the batch covariance structure alignment was determined based on the non-targeting control wells and applied to all wells in the screen. This was followed by another center scaling on the non-targeting control populations to ensure isotropic non-targeting control distributions.

### Training and evaluating elastic-net classifiers for gene knockdown phenotype assessment

For each dual guide construct, all the associated single-cell Plexus embeddings were collated and split into a 50-50 train-test split, such that all cells from the same well were in the same split. This split was also performed for the non-targeting control guides. This ensured that the evaluation on the test set was a proper test of the generalizability of the phenotype and not a function of overfitting to any well-specific phenotypes. The elastic net models were then fit on the train split with an equal weighting of the L1 and L2 norms. AUROCs were calculated on the predictions for the test set to evaluate the model’s performance and, hence, the magnitude of a generalizable phenotype.

### OPLS model

All Orthogonal Partial Least Squares models were fit following the approach described by Trygg et al.^38^ We performed Orthogonal Partial Least Squares (OPLS) analysis to identify latent components that could distinguish between different phenotypic states using Plexus embedding features. The OPLS model was trained on non-targeting control samples to separate the phenotypic variation associated with Tau mutation status or gene perturbation from cell-to-cell heterogeneity orthogonal to this variation. To fit the OPLS model, we first centered and scaled the predictor matrix *X* (Plexus embeddings) and the response vector *y* (Tau mutation status or gene perturbation status) to have zero mean and unit variance

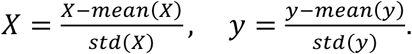

We then identified a weight vector *w* that maximized the covariance between *X_w_* and *y* by solving the following optimization problem

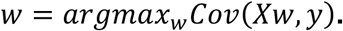

Using *w* we then computed the predictive component or the score *t*, which we term the disease-axis or predictive-axis in the case of the tau mutation status or the gene perturbation status, respectively.

*t = X_w_*.

The loading vector *p* was calculated to describe the relationship between *t* and the original data *X*.

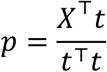

To calculate the orthogonal component, we then deflated *X* to remove the predictive variation, leaving us with *X*_o_.

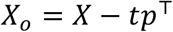

Using *X*_o_ we then calculated *w*_o_ by removing the components of the predictive loadings *p* that are aligned with *w*.

We then calculated the orthogonal component or ‘off-axis’ score as

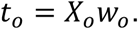

For the Tau mutation phenotype analysis, we performed the following statistical screening methods described below. We projected the Plexus embeddings from gene-targeting conditions onto both the disease axis and the orthogonal axis. To quantify the shift caused by each gene knockdown, we calculated the change in the distribution’s mean, or barycenter, along each axis.

The disease-axis score *S_disease_* for each gene knockdown gg was calculated as

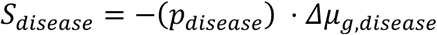

where *p_disease_* is the BH corrected p-value from a Mann-Whitney U test, and Δμ_g,*disease*_ represents the shift in the mean (barycenter) of the distribution along the disease axis compared to non-targeting controls.

The off-axis score *S_off_* was calculated as

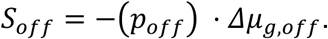

where *p_off_* is the corrected p-value for the orthogonal axis, and Δμ_g,*off*_ represents the shift in the mean of the distribution along the orthogonal axis. These scores allowed us to distinguish perturbations affecting the disease-relevant phenotype from those generating orthogonal variation.

### Interpretability analysis

To determine which manual features corresponded to the embedding features with the largest weights from the elastic net predictive axis. We defined the Plexus embeddings *H* ∈ *R*^n×d^_H_, the engineered feature array *F* ∈ *R*^n×d^_F_, and the elastic net predictive axis *w* ∈ *R*^d^_H_. The covariance matrix was calculated between all the Plexus embeddings and the engineered features Σ_HF_ = 1/n ^<^ *H*^T^*F*, and the elastic net predictive axis was projected into the manually engineered features space via the covariance matrix *w*_F_ = *Cov*(*H*, *F*)*w*. We then defined *w*_)_ as the discriminant projection score. This score was then used to determine the engineered features that most corresponded to the Plexus learned *KCNQ2* phenotype.

## Data Availability

The processed files containing the time series data, the model checkpoints, model embeddings in the form of h5ad files, and the raw dataset metadata files can be found at https://zenodo.org/records/14714574. Raw video files will be made available upon publication of this work.

## Code Availability

All code for plexus model training and inference is available at https://github.com/pgrosjean/plexus/tree/main (https://zenodo.org/records/15811302). All code for time series extraction from microscopy is available at https://github.com/pgrosjean/plexus-extract/tree/main (https://zenodo.org/records/15811338). All code for the multivariate Hawkes process simulation is available at https://github.com/pgrosjean/plexus-simulate/tree/main (https://zenodo.org/records/15811345).

## Biological Materials Availability

The cell lines generated in this study are available upon request and completion of a Material Transfer Agreement (MTA). Further information and requests for resources and reagents should be directed to and will be fulfilled by the Lead Contact, Martin Kampmann.

## Acknowledgments

We would like to thank the following members of the LGR for their technical support on the work presented in this manuscript: Phuong Nguyen, Scot Federman, Bryan Cunningham, and Brandon Kwan-Leong. We would also like to thank members of the GSK novel human genetics research unit for their helpful discussion. We acknowledge Duncan Muir, Noam Teyssier, Umair Khan, and Alex Lee for their scientific feedback while writing this manuscript.

This work is supported by the Laboratory for Genomics Research established by GSK, UCSF, and UC Berkeley and by grant DAF2018-191905 (https://doi.org/10.37921/550142lkcjzw) from the Chan Zuckerberg Initiative DAF, an advised fund of the Silicon Valley Community Foundation (funder https://doi.org/10.13039/100014989) (M.J.K.)

## Extended Data

### Extended Data Tables

**Extended Data Table 1:** Simulation parameters for the eight distinct neuronal activity phenotype classes used for linear probing.

| Phenotype Number | $\alpha$ | $\beta$ | $\gamma$ | $S_\beta$ | $S_\gamma$ | $\mu_{mean}$ | $\mu_{std}$ | Graph connectivity |
| --- | --- | --- | --- | --- | --- | --- | --- | --- |
| 1 | 8.3e-2 | 1.6 | 0.2 | 175 | 500 | 0.05 | 0.2 | 1.0 |
| 2 | 8.3e-2 | 2.0 | 0.2 | 25 | 500 | 0.05 | 0.2 | 1.0 |
| 3 | 0 | 2.0 | 0.2 | 25 | 500 | 2 | 0.2 | 0.2 |
| 4 | 0 | 2.0 | 0.2 | 25 | 500 | 0.2 | 0.2 | 0.2 |
| 5 | 8.3e-2 | 2.0 | 0.2 | 25 | 500 | 0.3 | 0.2 | 0.7 |
| 6 | 1.7e-2 | 2.0 | 0.2 | 25 | 500 | 2 | 0.2 | 0.7 |
| 7 | 8.3e-3 | 4.0 | 0.3 | 100 | 250 | 8 | 3 | 0.3 |
| 8 | 1.5e-2 | 0.2 | 0.01 | 25 | 25 | 0.1 | 0.02 | 0.4 |

### Extended Data Figures

**Extended Data Figure 1:**
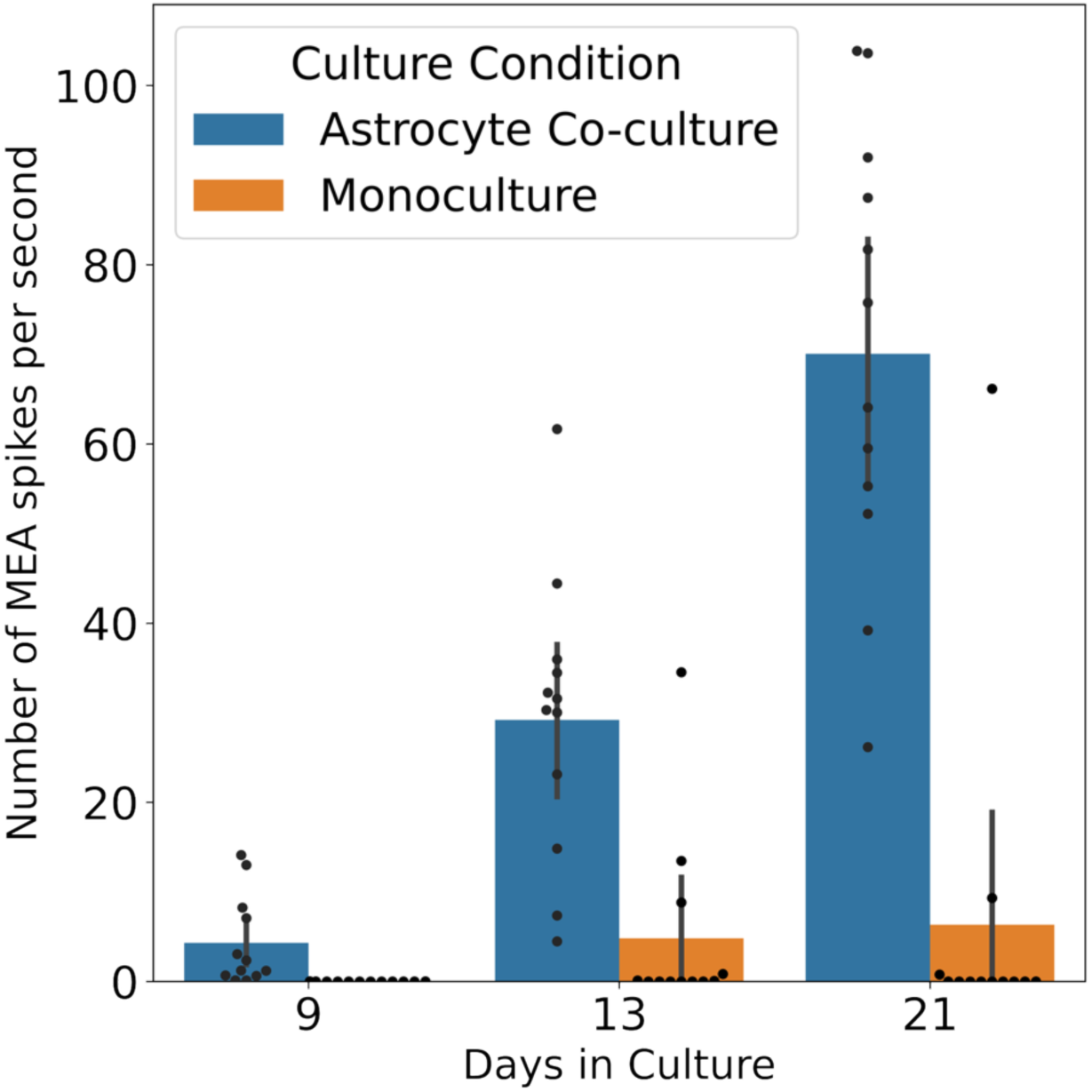
Activity measured by multi-electrode array in iNeuron monoculture vs astrocyte and iNeuron co-culture measured throughout culture.

**Extended Data Figure 2:**
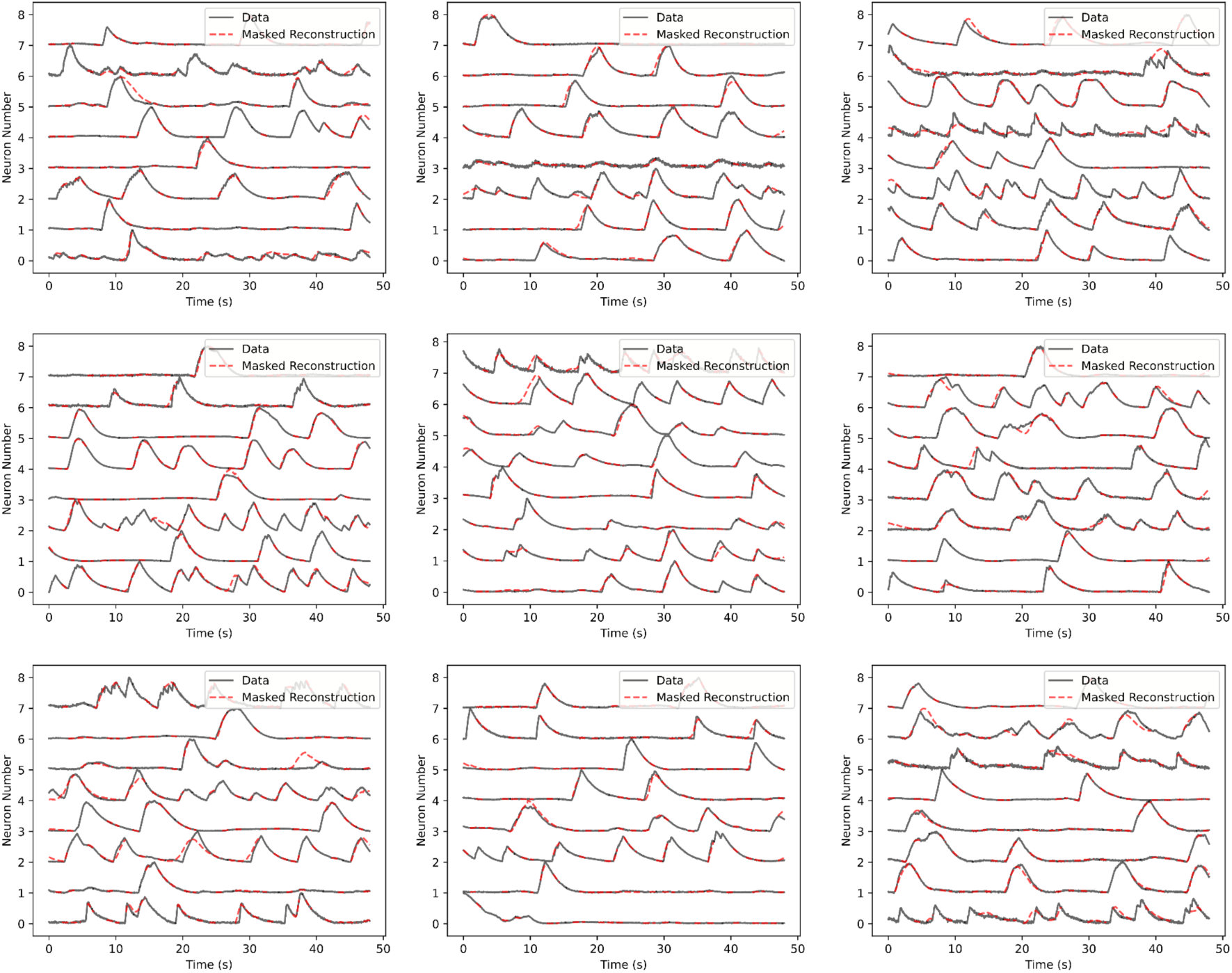
Plexus reconstructs masked neuronal activity dynamics. Nine representative examples of the reconstruction task with 50% masking. The original GCaMP6m data are shown in black, normalized between 0 and 1 for visualization purposes. The red traces depict the model’s reconstruction, specifically at the locations where masking was applied, highlighting Plexus’s ability to infer activity in masked regions.

**Extended Data Figure 3:**
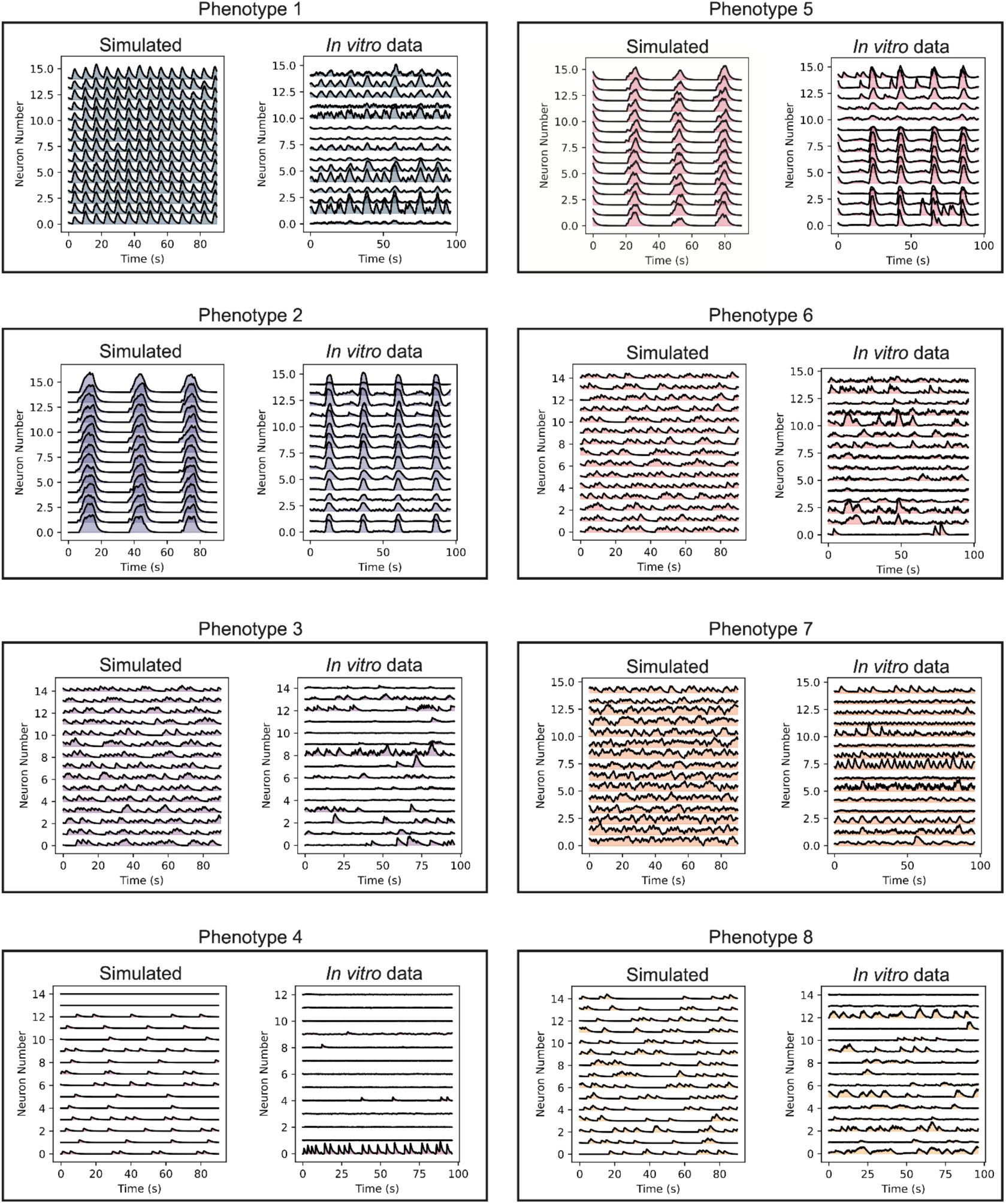
Representative traces of the eight simulated activity phenotypes (left traces) used for the linear probing classification task and the closest matching *in vitro* activity data (right traces) by distance in the Plexus embedding space.

**Extended Data Figure 4:**
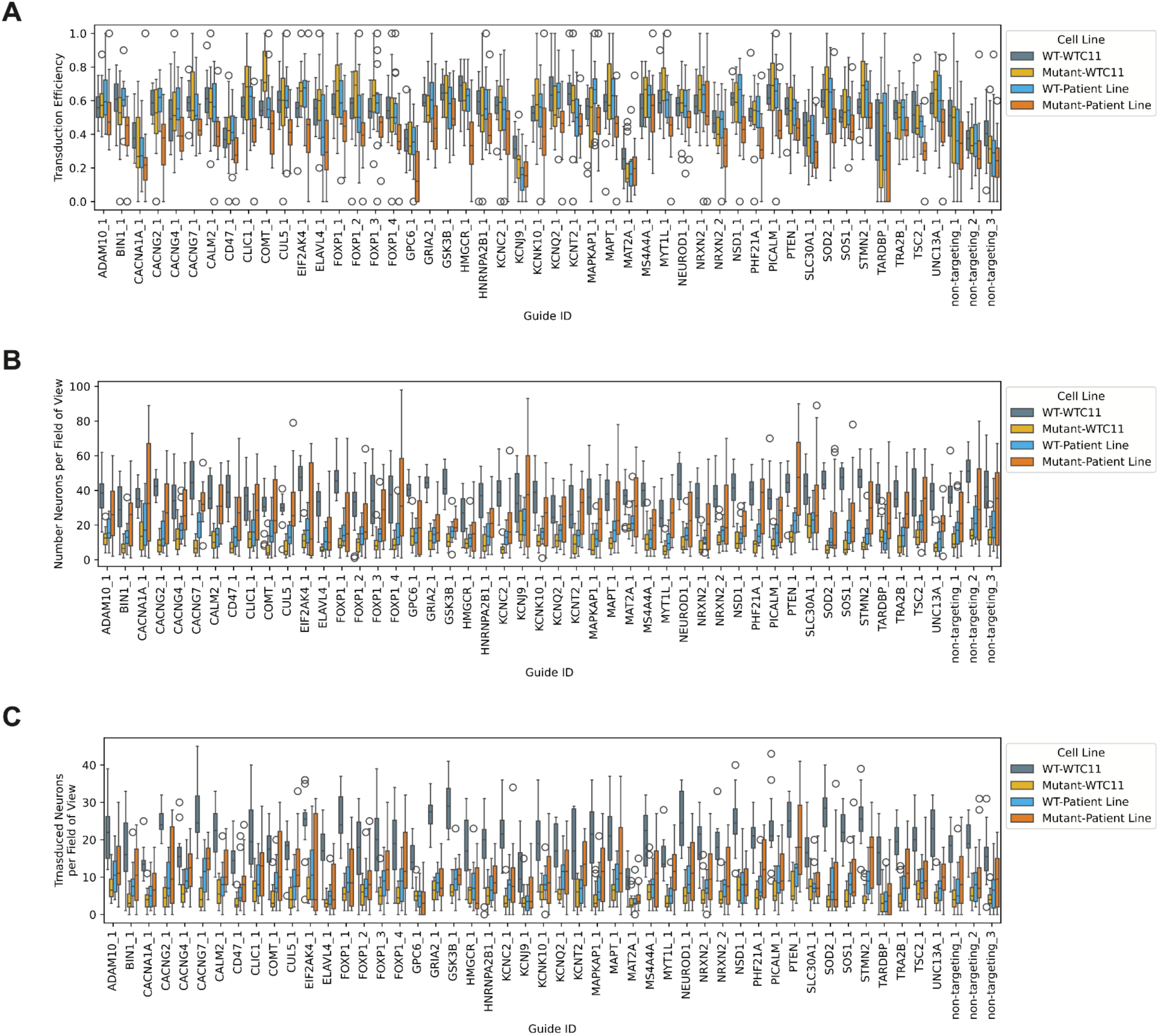
Neuron counts and transduction efficiency in the arrayed CRISPRi screen. (A) Boxplot (center line, median; box limits, upper and lower quartiles; whiskers, 1.5x interquartile range; points, outliers) of the transduction efficiency per well (n=32 for non-targeting guides, n=16 for all other guides). (B) The number of total neurons per field of view. (C) The number of transduced neurons per field of view.

