## Supplementary Tables for "Network-aware self-supervised learning enables high-content phenotypic screening for genetic modifiers of neuronal activity dynamics"

**Supplementary Table 1: CLS and register token ablation study for the neurostimulation experiment.**

| Plexus Linear Probe | AUROC (DMSO vs Neuroactive Treatment) | | | |
|---|---|---|---|---|
| | TeNT | TTX | 2 mM Ca2+ | 2 mM Mg2+ |
| Embedding – No Ablation | 1.00 +/- 0.00 | 1.00 +/- 0.00 | 1.00 +/- 0.00 | 1.00 +/- 0.00 |
| CLS Token | 0.99 +/- 0.00 | 1.00 +/- 0.00 | 1.00 +/- 0.00 | 1.00 +/- 0.00 |
| Register Token 1 | 0.99 +/- 0.00 | 1.00 +/- 0.00 | 1.00 +/- 0.00 | 1.00 +/- 0.00 |
| Register Token 2 | 0.99 +/- 0.00 | 1.00 +/- 0.00 | 1.00 +/- 0.00 | 1.00 +/- 0.00 |
| Register Token 3 | 0.98 +/- 0.01 | 1.00 +/- 0.00 | 1.00 +/- 0.00 | 1.00 +/- 0.00 |
| Register Token 4 | 0.99 +/- 0.01 | 1.00 +/- 0.00 | 1.00 +/- 0.00 | 1.00 +/- 0.00 |
| Register Token 5 | 0.99 +/- 0.00 | 1.00 +/- 0.00 | 1.00 +/- 0.00 | 1.00 +/- 0.00 |
| Embeddings – CLS/Register Ablation | 0.96 +/- 0.02 | 1.00 +/- 0.00 | 1.00 +/- 0.00 | 1.00 +/- 0.00 |

**Supplementary Table 2: Dual guide RNA CRISPRi library**

| Target Gene | PS1 | PS2 |
|---|---|---|
| STMN2 | GAAGGGTCCGGCTACAGCAG | GGTGGCGAAGGCAAAGGGTC |
| PHF21A | GCCTGGGCGGAGGGATCCCC | GGCTGGCTGGCTGTCCCCCC |
| MAPT | GGATTCGCGCAGACCCCAGG | GGGCTGCAGTCGAGAGTGAA |
| GPC6 | GTTGGGGATATACGAGATCT | GCAAAGGTGGGAGCGCGCGC |
| ATP13A3 | GGGCAGGGCGAGAACAAGGG | GATCACGCTGCGGGGAGAGC |
| UNC13A | GGAACCAAGATGGCCGGTGG | GCCCGGCGTGAGCCAAGCGC |
| ADAM10 | GGCGTTGCCGGCCCCTGAAG | GCATCCCCGCCGCCCAGGAG |
| CACNA1A | GCCGGCAGCCTCAGCATCAG | GTGTCCCGAGCTGCTATCCC |
| CLIC1 | GGGGATTTGATTGGCGCAGT | GGGTGTTTCAAATAAATAAG |
| L2HGDH | GGAAGCCACTTGACCCTCCA | GCTTCTGAGCGCTGAGGGAG |

| | | |
|---|---|---|
| STX1B | GCGGCACAGGGTAGGATGGA | GTGCGGCTGCGGCACAGGGT |
| KIF5A | GAGAGACTAAGGCTCTGCGG | GGCGCGCTGTCTCTCTCCTG |
| SLC30A1 | GTGAGGAGCTAGAGAGGCGG | GTGCTTGCTGGTCTCCTCTG |
| TARDBP | GGGAAGTCAGCCGTGAGACC | GCACCCGCTAGGCCGCTGCT |
| BIN1 | GCACGCCGCGCACCCGACAG | GCAGCGGAGCCAACTGACGG |
| MAT2A | GGAGCGAACGAAGCAGCGGG | GCGCGGAGCGAACGAAGCAG |
| MYT1L | GGTGCTTCAACAAGACTGCA | GACAATGAGCAAAAAACCCG |
| TRA2B | GAGCCCGTGCGGAGGCGGTG | GGTTAGAGCCCGTGCGGAGG |
| EIF2AK4 | GCAGCGCTGCGCCCAAGGCA | GGCCCACCGCCGCCCAGGCA |
| MAPKAP1_1 | GTGAAGCTCTGGGGGAAGAC | GCCTAGGGTGAAGTAGATCG |
| CALM2 | GCGGTGGAGCGGCCCCTGAG | GTGGATGCGGCGGAGGGATC |
| PTEN | GGTTAGAAAAGACGAAGAGG | GCGCCTGTGAGCAGCCGCGG |
| MAPKAP1_2 | GTGTGCGGCTCGGGGTAATA | GGGGGCATGAGGGCTAACCC |
| FOXP1_1 | GGCAGAAGCGAGAGTTCCGT | GGAGCCCTCAGATGAAGTGA |
| FOXP1_2 | GACCCCGCTGGGCACTCCAG | GCCGGGAAGCTAGTCCGGAG |
| TREM2 | GGTAAGATGAGCAGCCGGAG | GAGGAGGGTGTGAAGAATAT |
| HNRNPA2B1 | GGCGGCGGCAGCGGCTCTAG | GGCCCCGCTGGGGTGAAAGT |
| NRXN2_1 | GCTTCTGTCGGAGCCCGCGG | GGATGCCGGCTTCCCTCAGG |
| KCNC2 | GACCCCAGCGCCCAGGGAAG | GGGCTCAGGAAAGAGTGGGT |
| ELAVL4 | GATCTACATCCTAGAATCGG | GTTGTGTAGAGAGTGCGGGT |
| FOXP1_3 | GCCGGGAAGCTAGTCCGGAG | GGGAGCCCAGCCAGCGCCGG |
| GIGYF2 | GGCAGGGGAGCGACACGGAA | GCGGAACGGCTGGTTACCTG |
| KCNK10 | GTGTGCGCACACGCCTTCGG | GTAGGGGCTGCGCAGCCTGA |

| | | |
|---|---|---|
| MS4A4A | GGCCATCAGCCCGAAAGCCT | GTGTAGGTCTGATGCATGCA |
| NEUROD1 | GCCCGCGCGGCCACGACACG | GCAGGAGGCGCGGCGTCCGG |
| CD47 | GCGGTCGGTCCTGCCTGTAA | GTCGGTCCTGCCTGTAACGG |
| NRXN2_2 | GGATGCTCCGCGAGTCAGGG | GGAGGATGCTCCGCGAGTCA |
| CACNG7 | GGAAGGGGCGCCCCTCGAGG | GCGGCTCCGTCCGCCGAGTG |
| TSC2 | GGGGAACGCAGGGCCGCACG | GTTGGGGCGAAAGGGGGCAG |
| SOS1 | GTCCAGCGCTACACCGGCGG | GGGAAGCTCCAGCGCTACAC |
| CACNG2 | GCCAGCCGGTAGCGAGTGGG | GCACCCGTAATGCAGTAGGT |
| KCNQ2 | GCCCAAGCCCGGCAGGAGTG | GTGCAGAAGTCGCGCAACGG |
| SOD2 | GGGCGCAGGAGCGGCACTCG | GAGCGGCTTCAGCAGATCGG |
| NSD1 | GGCTGCCGCGAACTTCCTCC | GGCGGCGGCGGAATAGGCCG |
| PICALM | GTCGGCTTCACTCACAGTCC | GCTCGTGTCACCCGCGGAGT |
| CACNG4 | GCCGGAGCGCAGGCACGCCC | GCGCTGGGGCTCAAACTCCG |
| CUL5 | GCCCAAAGAGCCGAACCCGG | GGCGGCGGGTGCAACCACAA |
| KCNT2 | GTATTGCGGAGTACAGGGAG | GCTGTGGCCGAGAGAGGGAT |
| KCNJ9 | GCCCCCACGGGCCCCCCGAA | GCACGGGCCCCCCGAAGGGT |
| GSK3B | GGGCCCGGCGAACTAGAGGG | GGGAGTCGCGAGTCAGTCAG |
| FOXP1_4 | GGGCGACTCCTGCTGCACAG | GCAGGTGGCTAACCAGGGCT |
| COMT | GCATCGTCGTGGGGCTTCTG | GTCCGGCGCAGGCAGCGCGG |
| HMGCR | GACTAGAGGCCACCGAACCC | GCACCTCCAGATCTCACTAG |
| GRIA2 | GCCCCGCTCCAGAAGTCCGG | GCTCCAGAAGTCCGGAGGAC |
| SCN1A | GTGAAGAAGTTGAAGCTGTC | GTTGGTGCTACAACAGTCCA |
| non-targeting_00999 | GCTTGAATGGCTCCACCTAA | GAAAGGAGCCCCTGCAACCG |

| | | |
|---|---|---|
| **non-targeting_02396** | **GAATCGCCCAGCCAAGGCGC** | **GCTCCGGCGACGACAGGCGC** |
| **non-targeting_02496** | **GGGCGTCTACCAAGCCTCAT** | **GGCGGATAACGCATATGAAC** |
| **non-targeting_02443** | **GTTAAGAGCTCAGACGGCGG** | **GACGGCGCGTGCGCTTAGTG** |
| **non-targeting_01460** | **GAGGCGCAAGCGGACGTGTG** | **GGCCGCACGATAGTGTCCTA** |

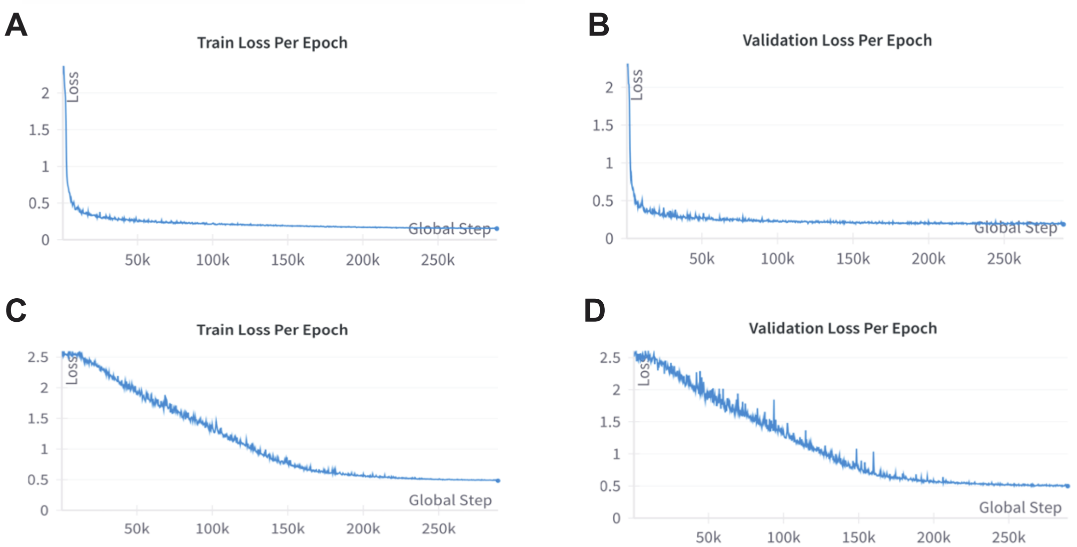


**Supplementary Figure 1**: **Loss curves for Plexus model training for CRISPRi dataset.** (A) Training loss curve for Plexus using the Plexus cell identity embeddings (learnable rank order set-invariant discrete tokens). (B) Validation loss curve for Plexus using the Plexus cell identity embeddings (learnable rank order set-invariant discrete tokens). (C) Training loss curve for Plexus using mean signal cell identity embeddings (mean signal as set-invariant continuous tokens). (D) Validation loss curve for Plexus using mean signal cell identity embeddings (mean signal as set-invariant continuous tokens).

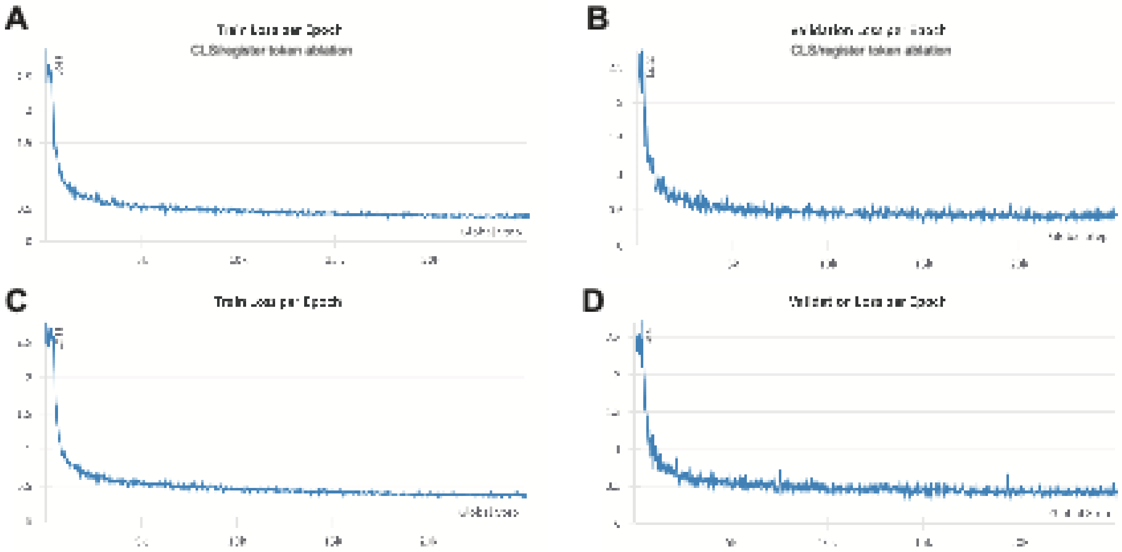


**Supplementary Figure 2: Loss curves for training with and without CLS and register tokens.** (A) Training loss curve for Plexus without CLS or register tokens. (B) Validation loss curve for Plexus without CLS or register tokens. (C) Training loss curve for Plexus with CLS or register tokens. (D) Validation loss curve for Plexus with CLS or register tokens.

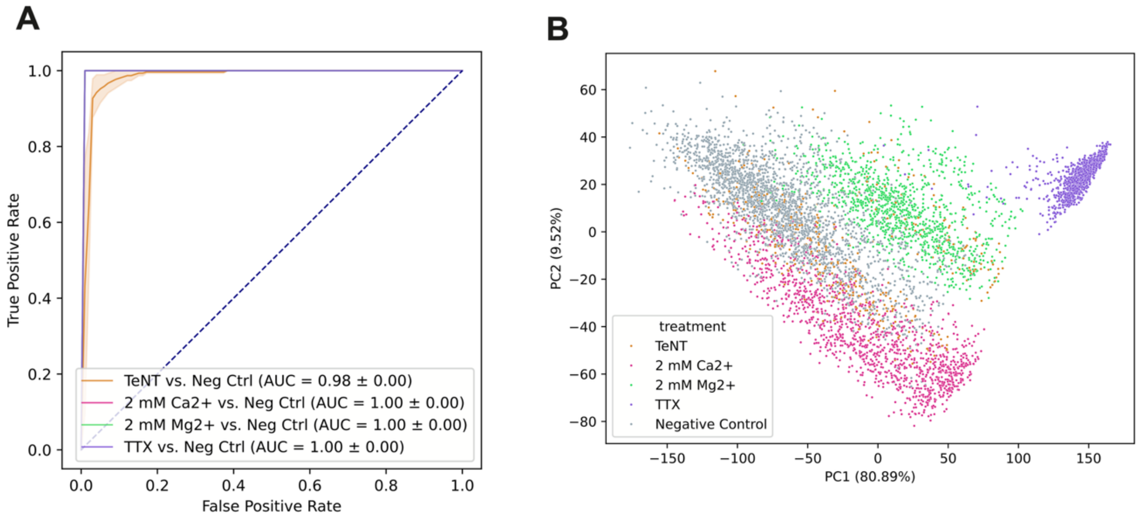


**Supplementary Figure 3**: **CRISPRi dataset trained Plexus out-of-distribution probing on the neuroactive stimulation dataset.** (A) ROC curves for the Plexus model trained on the CRISPRi dataset used for linear probing for neuroactive stimulation. (B) PCA plot of the neuroactive stimulation activity embeddings from the CRISPRi dataset-trained Plexus model, colored by neuroactive stimulation.

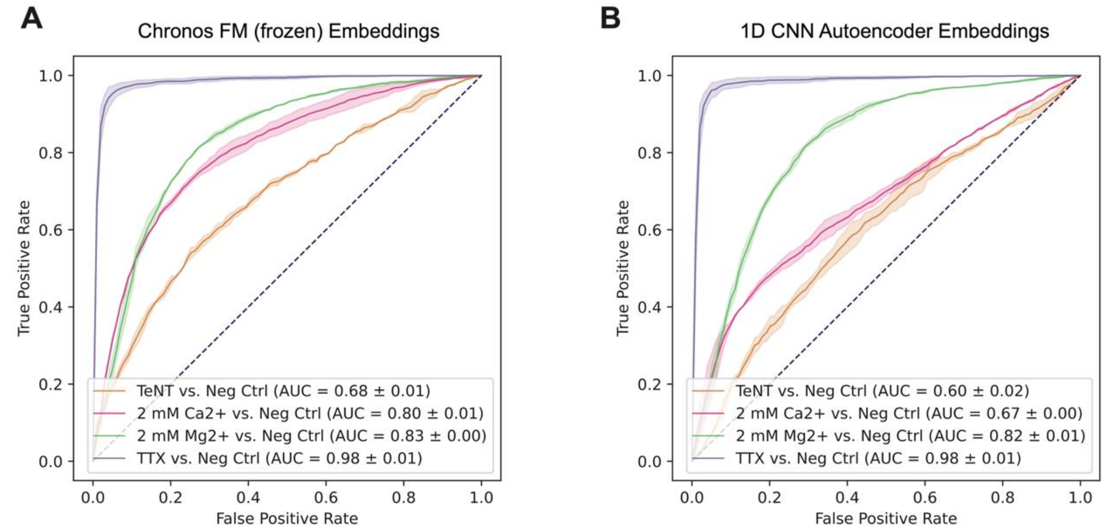


**Supplementary Figure 4**: **ROC curves for additional representation learning model baselines for the neurostimulation binary classification tasks.** (A) ROC curves for linear probing of the frozen Chronos Foundation Model. (B) ROC curves for the 1D CNN autoencoder baseline.

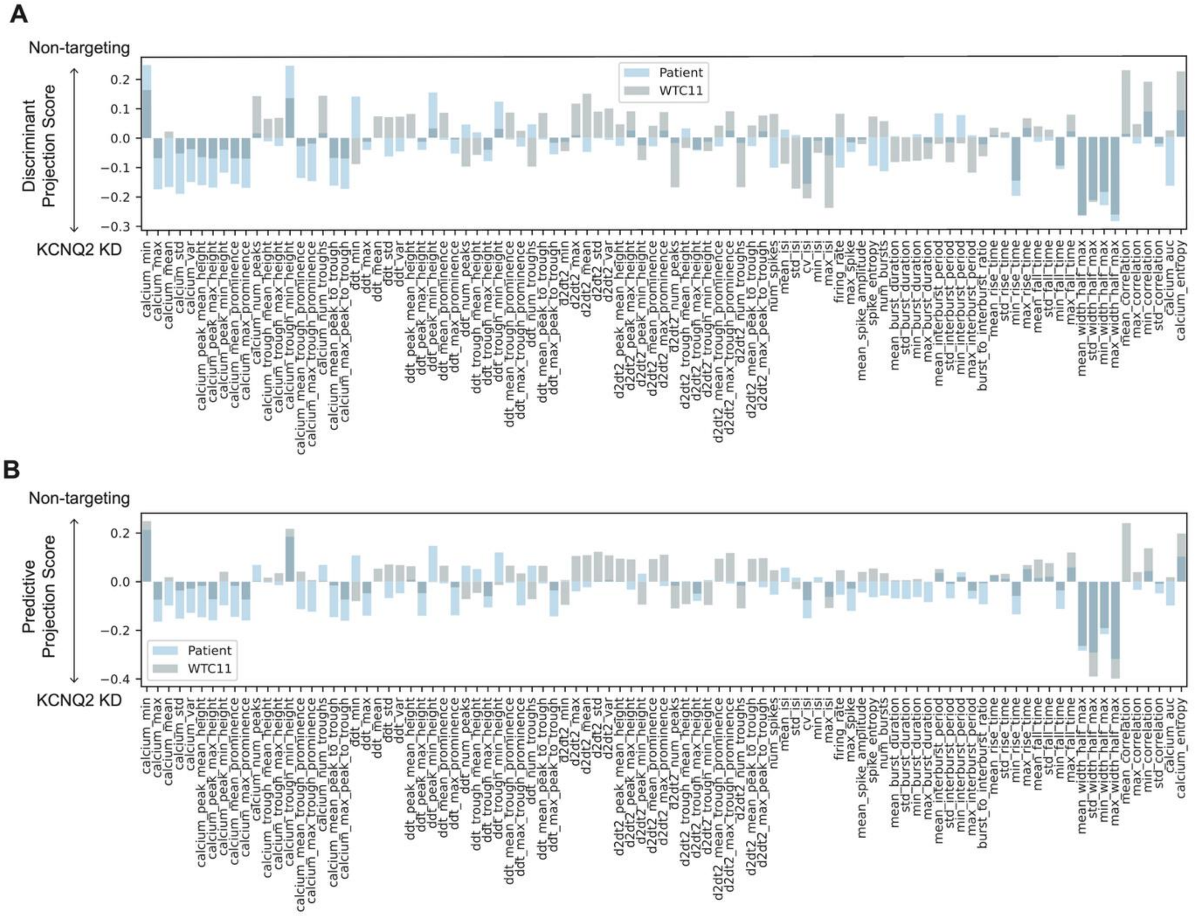


**Supplementary Figure 5**: **KCNQ2 knockdown induces a distinct phenotype in two iPSC-derived iNeuron cell lines determined by LDA and OPLS.** (A) The mean Linear Discriminant Analysis (LDA) axes from both cell line models are projected onto the covariance matrix of Plexus embedding features and engineered features. The resulting discriminant projection scores reflect feature influence: negative scores indicate a KCNQ2 knockdown-like phenotype, while positive scores align with non-targeting controls. (B) The mean OPLS predictive axes from both cell line models are projected onto the covariance matrix of Plexus embedding features and engineered features. The resulting discriminant projection scores reflect feature influence: negative scores indicate a KCNQ2 knockdown-like phenotype, while positive scores align with non-targeting controls.

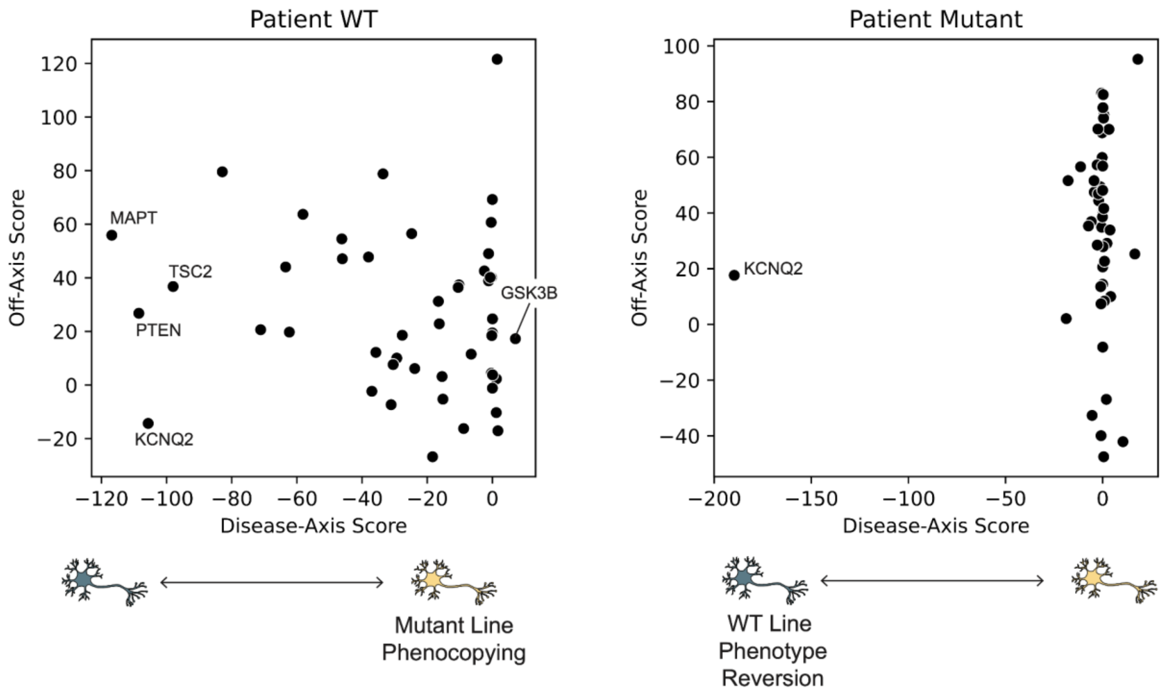


**Supplementary Figure 6**: **Patient line gene knockdown OPLS projections.** (Left) Patient wild-type Tau line disease-axis vs off-axis scatter plot demonstrates that most gene knockdowns in this cell line seem to move the cells away from a mutant-like phenotype. (Right) Patient mutant Tau line disease-axis vs off-axis scatter shows that most gene perturbations drive off-predictive-axis effects.

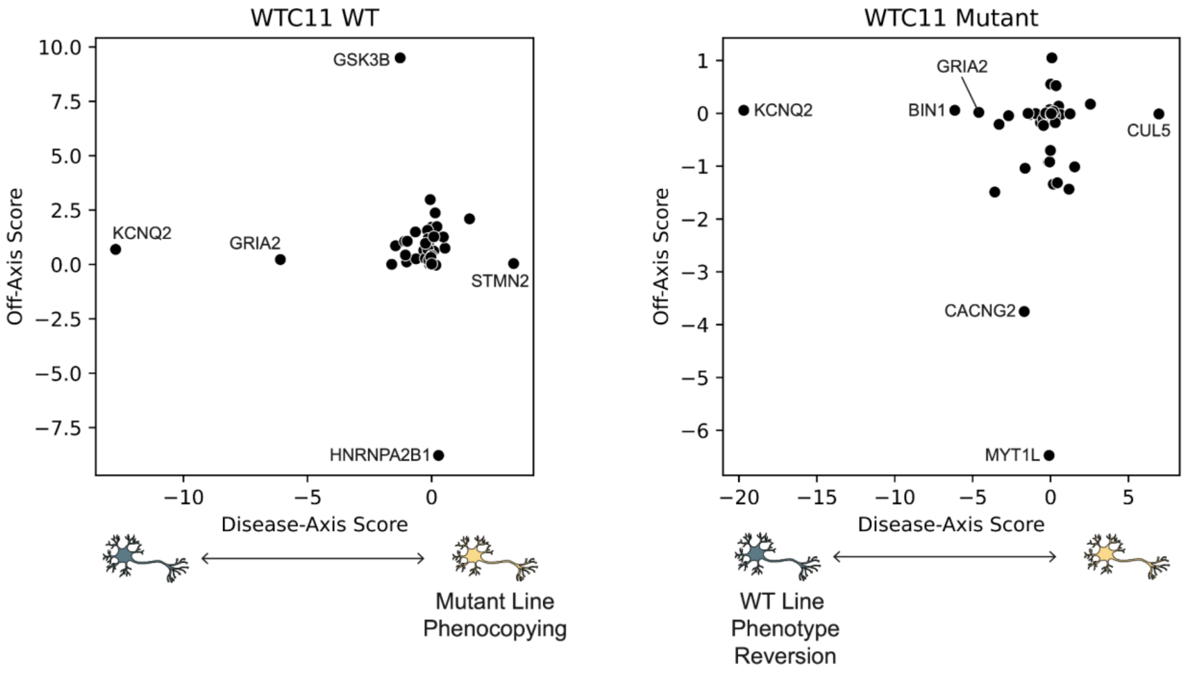


**Supplementary Figure 7**: **Manual Features WTC11 gene knockdown OPLS projections**. (Left) WTC11 wild-type Tau line disease-axis vs off-axis scatter plot from OPLS model using signal processing-based manual features. (Right) WTC11 mutant Tau line disease-axis vs off-axis for manual features.
